# Peptide location fingerprinting identifies species- and tissue-conserved structural remodelling of proteins as a consequence of ageing and disease

**DOI:** 10.1101/2022.01.31.478441

**Authors:** Alexander Eckersley, Matiss Ozols, Peikai Chen, Vivian Tam, Liam J. Ward, Judith A. Hoyland, Andrew Trafford, Xi-Ming Yuan, Herbert B. Schiller, Danny Chan, Michael J. Sherratt

## Abstract

Extracellular matrix (ECM) in the intervertebral disc (IVD), lung and artery are thought to undergo the age-dependant accumulation of damage by chronic exposure to mechanisms such as reactive oxygen species, proteases and glycation. It is unknown whether this damage accumulation is species-dependant (via differing lifespans and hence cumulative exposures) or whether it can influence the progression of age-related diseases such as atherosclerosis. Peptide location fingerprinting (PLF) is a new proteomic analysis method, capable of the non-targeted identification of structure-associated changes within proteins. Here we applied PLF to publicly available ageing human IVD (outer annulus fibrosus), ageing mouse lung and human arterial atherosclerosis datasets and identified novel target proteins alongside common age-associated differences within protein structures which were conserved between tissue regions, organs, sexes and species and in age-related disease. We identify peptide yield differences across protein structures which coincide with biological regions, potentially reflecting the functional consequences of ageing or atherosclerosis for macromolecular assemblies (collagen VI and fibrin), enzyme/inhibitor activity (cathepsin B and alpha-2 macroglobulin), activation states (complement C3 and thrombin) and interaction states (laminins, perlecan, fibronectin, filamin-A, collagen XIV and apolipoprotein-B). Furthermore, we show that alpha-2 macroglobulin, prothrombin, collagen XIV and apolipoprotein-B all exhibit possible shared structural consequences in IVD ageing and arterial atherosclerosis, providing novel links between an age-related disease and intrinsic ageing. Crucially, we also demonstrate that fibronectin, laminin beta chains and filamin-A all exhibit conserved age-associated structural differences between mouse lung and human IVD, providing evidence that ECM, and their associating proteins, may be subjected to potentially similar mechanisms or consequences of ageing across species, irrespective of differences in lifespan and tissue function.

## Introduction

In contrast to the dynamic intracellular environment where proteins are replaced hourly or daily, extracellular matrix (ECM)-rich tissues are susceptible to a unique form of ageing which is governed by the longevity of the constituent extracellular proteins. For instance, collagens in the intervertebral disc (IVD) [1] and elastic fibres in lung [2] remain present throughout the human lifespan. As a consequence of this extremely low turnover, many ECM components in tissues such as lung, IVD and artery are thought to undergo the age-dependant accumulation of damage via chronic exposure to a variety of mechanisms. These include reactive oxygen species (ROS), markers of which are known to be upregulated in aged IVD [3, 4], artery [5] and lung [6]. ROS can directly oxidise sensitive amino acid residues (Cys, Met, Trp, Tyr and His) leading to changes in higher order structure [7, 8]. Damage to ECM components can also be incurred by proteolytic degradation and fragmentation through chronic exposure to proteases [9, 10], which are often elevated in aged and degenerate tissues; as seen for matrix metalloproteinases (MMPs) 1 and 2 in both aged IVD [11] and artery [12]. Glycation of proteins and cross-linking of long-lived ECM components by advanced glycation end products (AGEs) in ageing connective tissues like IVD [13], lung [14] and artery [15] can also affect the stiffness, accessibility and interactivity of matrix proteins with age. Although the mechanical forces involved may differ between these tissues, repeated mechanical deformation can also lead to ECM fragmentation [9,16,17].

Despite evidence of these ECM-specific ageing mechanisms in the connective tissues of human and model organisms, it is unknown whether their effects are species-dependant, because of differing lifespans. For instance, with an approximate life expectancy of only 2.5 years, mice have long been used as a biological model for human ageing [18]. Therefore, the determination of whether the matrisome ages comparably within these species is a relevant question in ageing research. Furthermore, it is also unknown whether the gradual accumulation of damage to long-lived ECM components can influence the progression of age-related diseases, some of which, such as arterial atherosclerosis and lung chronic obstructive pulmonary disease (COPD), have been attributed to an accelerated form of tissue ageing [19, 20]. Therefore, the identification age-susceptible proteins that are conserved between species, organs and with age related diseases, and the characterisation of shared mechanisms and functional consequences, is crucial for the understanding of connective tissue ageing and the discovery of new therapeutic targets.

The age-dependant degeneration of ECM architectures, such as for elastic and collagen fibres, is well documented in tissues such as the IVD, lung, aorta and skin [21–24]. However, time-dependent changes to higher order molecular structure, and their downstream consequences, are challenging to identify as they may be independent of changes in protein transcription, abundance, or architecture. Liquid chromatography tandem mass spectrometry (LC-MS/MS) has been used previously to investigate targeted post-translational modifications to protein structures [25]. However, due to the irregular and often non-specific nature of age-associated, variable modifications, accrued over time, this approach is ill-suited for the protein structure-related study of ageing and disease in ECM-rich connective tissues.

Peptide location fingerprinting (PLF) is a new and emerging proteomic analysis method, capable of the non-targeted and unbiased identification of structure-associated changes within proteins by mapping and quantifying LC-MS/MS-detected tryptic peptides within specific protein regions [26–28]. This approach takes advantage of differences in the regional digestibility of proteins, because of solubility, stability and enzyme susceptibility, to measure characteristic peptide patterns (fingerprints) across their higher order structures. Therefore, by comparing LC-MS/MS datasets, PLF can robustly identify significant fluctuations in peptide yield patterns across protein structure as a consequence of ageing [26], induced damage [28] and even tissue diversity [29]. These identified changes in structure can be reflective of damage modifications or changes in protein conformation, interaction states, activation or synthesis. Due to their insolubility (resultant from highly cross-linked networks), PLF is particularly effective at measuring structure-associated changes within ECM assemblies. For instance, we have previously used PLF to reliably identify ultraviolet radiation-induced, regional damage within ECM proteins as an *in vitro* model of skin photoageing [28], as well as tissue-specific differences between eye- (ciliary body) and skin-derived fibrillin-1 [29].

Recently, through the development of the Manchester PLF webtool (MPLF), we applied PLF as a proteomic biomarker discovery tool to both self-generated human LC-MS/MS datasets of skin photoageing [26] and to historical tendon [26] and IVD [27] ageing datasets, sourced from a public database (PRIDE). In these studies, we identified potential photoageing- and ageing-affected proteins which were independent of changes in whole protein abundance, demonstrating that PLF identifies structure-associated differences that are unique to the methodology. Most crucially however, we also showed that regional peptide yield differences within these proteins could be correlated to specific biological mechanisms, such as increased collagen I, II and V synthesis in young IVD compared to aged (due to the higher presence of peptides from C-terminal propeptide regions), and increased collagen I and II degradation in aged IVD compared to young (due to the higher presence of peptides from regions downstream of their prominent MMP cleavage sites) [27].

It is clear that PLF can be used as a proteomic screening tool to interrogate structure-associated changes to ageing ECM proteins in multiple tissues, using historical label-free LC-MS/MS datasets [26, 27]. In this study, we aim to apply PLF to three previously published datasets representative of ECM-rich connective tissue ageing (mouse lung [30] and human IVD outer annulus fibrosus [31]) and age related-disease (human arterial atherosclerosis [32, 33]) **(Fig. 1)**. The IVD, lung and artery were chosen as they all undergo age-related remodelling [30,34–37] which can severely impact their function and in turn morbidity [38, 39] or mortality [40, 41] for the host organism. Although there are some differences in their respective ECM proteomes (matrisomes), many extracellular assemblies, including fibrillar collagens, elastic fibres and proteoglycans, are common to all three organs (and to other ECM-rich tissues). Although these ageing and atherosclerosis datasets were previously used to investigate global differences in proteome composition and specific differences in relative protein abundance, modification-related differences to ECM protein structures were not interrogated. This provides us with a unique opportunity to apply PLF not only to identify novel biomarker candidates, but also crucial evidence of age-associated differences to structure which are conserved between species, organs and in age-related disease.

**Figure 1.**
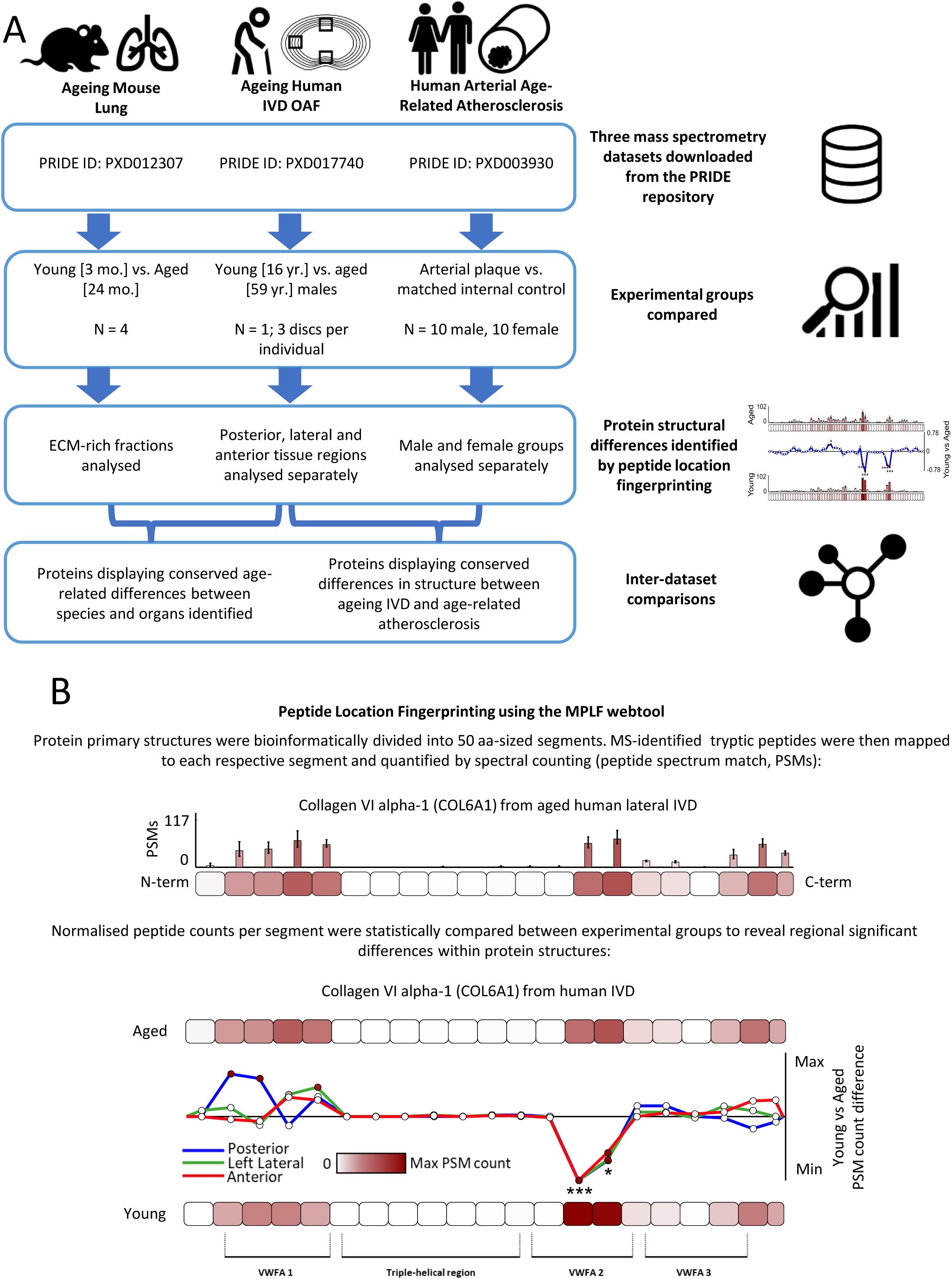
Experimental workflow and methodology for the application of PLF to ageing and age-related disease in three distinct tissues. Raw LC-MS/MS datasets derived from mouse lung [30], human IVD outer annulus fibrosus (AF) [31], and human atherosclerotic artery [32, 33] **(A)** were downloaded from the Proteomics Identification Database (PRIDE) repository. After peptide identification, PLF was used to compare peptide yield across protein structures between young and aged groups for mouse lung and human IVD and between atherosclerotic plaque and internal control artery to reveal proteins with age- or age-related disease-dependant structural differences. Additionally, these analyses were stratified by tissue location for IVD (posterior, left lateral and anterior) and by sex for artery to expose region- and sex-related similarities and differences. Finally, peptide yield differences across affected proteins, commonly identified between datasets, were cross compared to reveal matching patterns of ageing and disease and to expose evidence of conserved mechanisms of change in key proteins. In PLF analysis **(B)** primary sequences of matched proteins were bioinformatically divided into 50 amino acid (aa)-sized segments. Collagen VI apha-1 is shown here as an example. Identified peptide sequences were then mapped and quantified within each segment by spectral count which were then summed. These were median normalised based on the experiment-wide total spectrum counts of their corresponding whole proteins which minimised the skew of whole protein relative abundance on structural comparisons. Modular fluctuations in peptide yield across structures were assessed by subtracting the average, normalised peptide counts per segment in one experimental group from the other and dividing them by the segment length (50 aa). Additionally, average, normalised peptide counts in each segment were statistically compared between experimental groups using Bonferroni-corrected, repeated measures ANOVAs (unpaired for young vs aged and paired for plaque vs control).

## Results and Discussion

### PLF reveals age-affected proteins in IVD outer annulus fibrosus exhibiting tissue region-conserved and -specific fluctuations in peptide yield within protein structures

The human ageing IVD dataset was previously used in the development of the spatiotemporal proteomic analysis platform: DIPPER (http://www.sbms.hku.hk/dclab/DIPPER/). This includes the detailed analysis of 11 spatially resolved tissue regions (posterior, lateral, anterior and central portions of inner and outer AF, and nucleus pulposus) of three discs (L3/4, L4/5 and L5/S1) sourced from two cadaveric males (aged 16 and 59 years respectively) [31]. This formerly published study revealed a profound shift in proteomic composition and abundance along the anteroposterior and lateral axes of aged disc, with the inner and outer AF regions converging. The inner AF portions of this dataset were previously analysed by PLF [27]; by comparing proteomes separately derived from the posterior, anterior and lateral regions, we were able to show tissue region-conserved and -specific differences in peptide yield patterns within ageing ECM proteins such as aggrecan and collagen I, II, and V alpha chains [27]. Here, we chose to focus on the outer AF (OAF), which previously showed a clear shift of ECM composition between aged and young [31], and is therefore most likely to harbour structure-associated evidence of ageing in long-lived matrix proteins. Since IVD ageing and degeneration is profoundly regional, we stratified this comparison between anterior, posterior and left lateral regions as before, to reveal tissue location-specific similarities and differences. Young and aged discs were compared: three vs three discs for lateral and anterior and three vs two discs for posterior (young posterior L4/5 dataset omitted due to insufficient peptide identification for PLF).

Peptides corresponding to 684 proteins in anterior, 800 in left lateral and 441 in posterior were identified in both young and aged disc OAFs by MS/MS ion searches (**Fig. S1**; peptide lists: **Tables S1 – S3**). Principal component analyses (PCAs) of peptide spectral counts demonstrated good separation of data between young and aged groups for all three tissue regions, with young discs clustering in both anterior and left lateral comparisons **(Fig. S2)**. PLF analysis led to the identification of 284 proteins in total (across all tissue regions) with protein regions exhibiting significantly different peptide yields between aged and young discs, 27 of which were shared between tissue regions **(Fig. S3)**. Classification analysis of these shortlisted proteins revealed ECM proteins as the major class, making up ∼44% of age-affected proteins which were shared between at least two tissue regions **(Fig. 2)**. These included collagens, elastic fibre proteins, basement membrane proteins and proteoglycans, indicative of extensive ECM remodelling within the OAF of aged discs compared to young.

**Figure 2.**
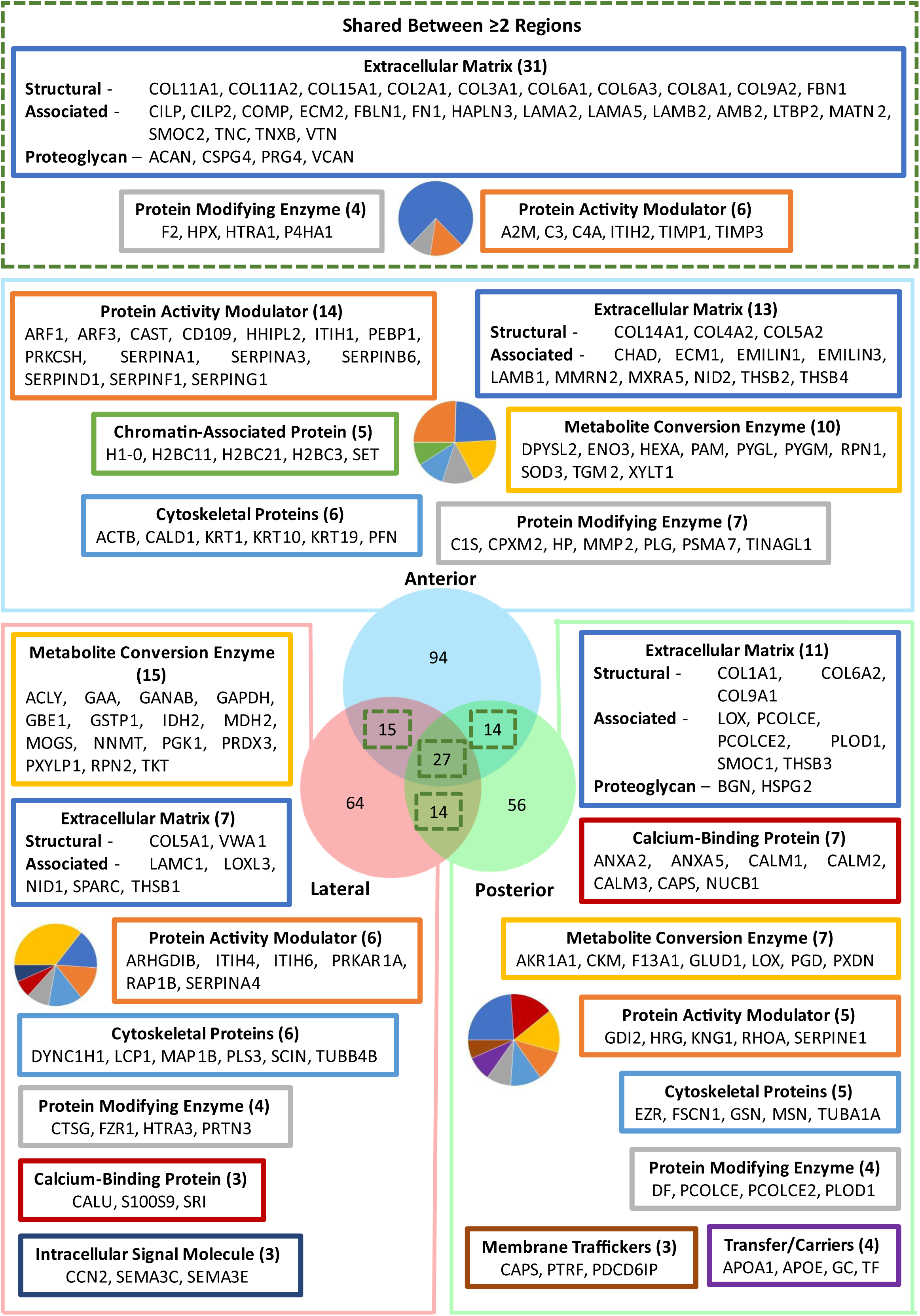
Classification analysis of proteins with structure-associated modifications revealed ECM proteins as the major class affected by age in IVD OAF. PLF analysis identified 150 proteins in anterior, 120 in left lateral and 111 in posterior tissue regions with significant differences in peptide yield across structure (22%, 15% and 25% of total proteins identified respectively per region; full PLF analysis results: **Tables S4 – S6**). PANTHER classification analysis [42] revealed ECM proteins as the major class of proteins affected in at least two tissue regions (classes and their protein identities are displayed with coloured frames matching their respective slices within the pie charts; minimum 3 proteins per class; total number of proteins per class indicated in brackets). Protein activity modulators (e.g. serpins), metabolite interconversion enzymes (e.g. superoxide dismutase, lysyl oxidase), protein modifying enzymes (e.g. MMP2, procollagen C-endopeptidase enhancers) and cytoskeletal proteins (e.g. keratins, tubulins) were also among the major classes structurally affected by age, but were predominantly tissue region-specific.

Peptide yield differences across the structures of PLF-identified proteins were further compared between posterior, lateral and anterior to investigate whether age-dependent fluctuations were consistent between IVD OAF regions. Additionally, by relating these differences to biological domains within the protein, it was possible to interrogate the functional consequences of these changes with age. Four ECM proteins in particular exhibited clear, profound tissue region-conserved differences in peptide yield between young and aged, with patterns that are indicative of unique consequences of ageing. These are the alpha-3 chain of the microfibrillar collagen VI (COL6A3), the proteoglycan versican and the ECM remodelling cartilage intermediate layer proteins (CILP)-1 and -2 **(Fig. 3)**.

**Figure 3.**
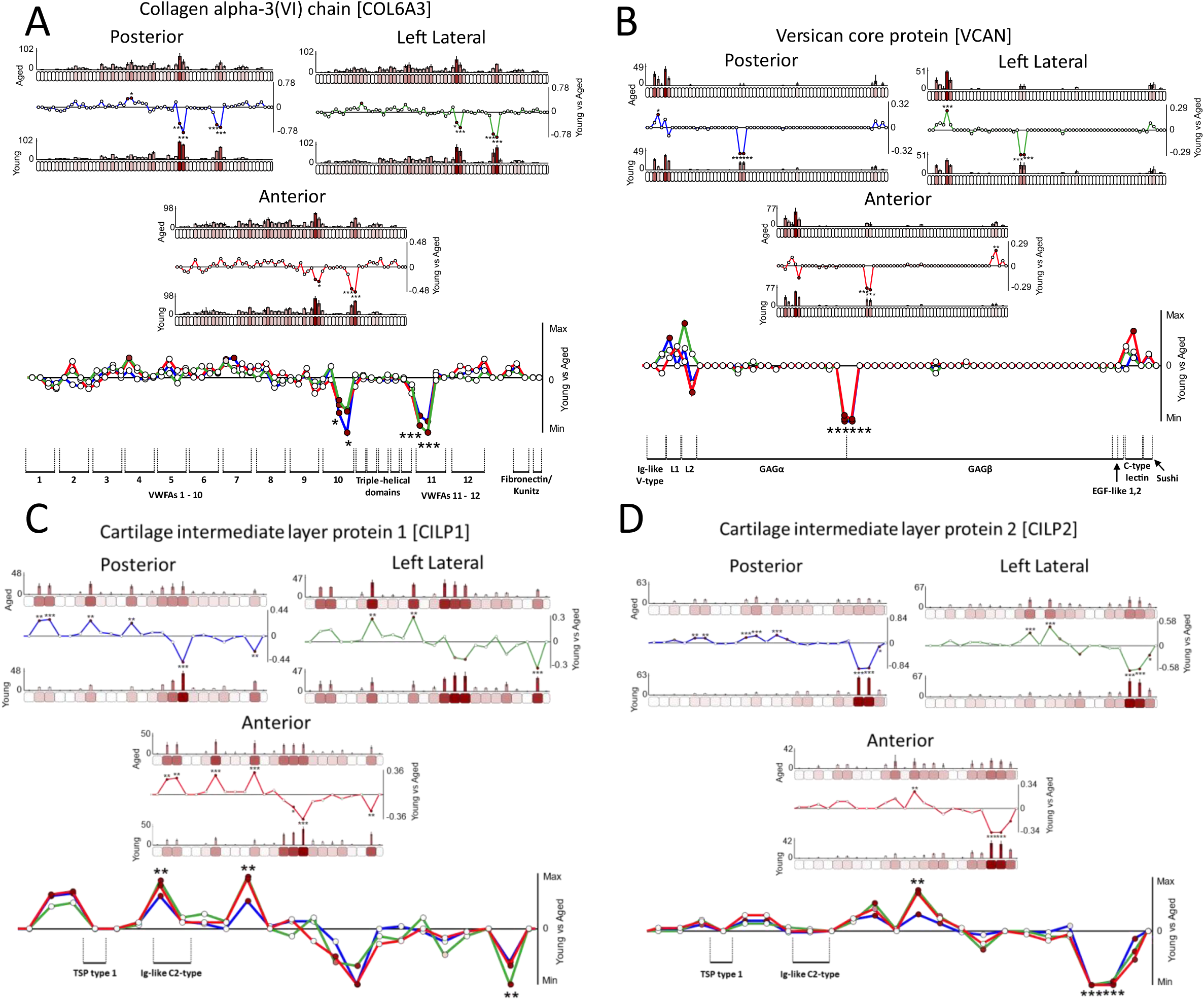
ECM proteins displaying tissue region-conserved differences in peptide yield across their modular structures between young and aged IVD OAF. Protein primary sequences were segmented into 50 aa-sized segments and LC-MS/MS-detected peptide sequences were quantified within each segment (bar graphs = average, normalised PSMs; error bars = standard deviation [SD]). Modular fluctuations in peptide yields across protein structures were assessed by subtracting the average, normalised peptide counts per segment in young from aged (Line graphs = aged-young PSMs/segment aa size [50]; below zero line = higher in young, above zero line = higher in aged) and statistically compared between groups (unpaired Bonferroni-corrected, repeated measures ANOVAs: *, p ≤ 0.05; **, p ≤ 0.01; ***, p ≤ 0.001; composite line graphs: stars = significant in all three tissue regions; aa ranges of Uniprot-sourced functional domains and regions are indicated). COL6A3 **(A)** contained multiple segments within the C-terminal half of the protein which displayed significantly higher peptide yields in young than in aged. These were consistent between posterior, lateral and anterior regions of the IVD OAF and coincided specifically with von Willebrand Factor type A (VWFA) domains 10 and 11. Versican **(B)** also contained two segments near the protein centre which also exhibited higher peptide yields in young than in aged and were consistent between all three tissue regions. These corresponded to the interface between glycosaminoglycan (GAG) α and GAGβ regions of the protein. The N-terminal half of CILP1 **(C)** yielded higher peptides in aged than in young (one significantly different segment coincided with the ig-like C2-type domain) whereas the C-terminal half yielded more peptide in young than aged. These peptide yield difference patterns were remarkably conserved between all three tissue regions, across the entire modular structure of CILP1. Similarly, this pattern was also tissue-region conserved for CILP2 **(D),** which contained a segment at the protein centre which yielded significantly higher peptides in aged than in young and two near the C-terminal end which yielded significantly more in young than in aged.

Collagen VI exists as a multifunctional microfibril composed of trimeric alpha chains (mainly alphas-1,-2 and -3) that regulates tissue homeostasis [43] through its extended network with other ECM components [44] and cell interactions [45]. Previous analysis of this dataset showed that alpha chains-1,-2 and -3 all had significantly lower relative abundances in aged IVD OAF compared to young, indicating a reduction of collagen VI with age [31]. Here, we observed two specific regions of COL6A3 which displayed significantly higher peptide yields in posterior, lateral and anterior tissue regions of young OAF than in aged. These corresponded directly to the VWFA domains 10 and 11 **(Fig. 3A)**, which are located within the globular double beaded region of the collagen VI microfibril and directly flank the inter-bead triple helical region [43]. Since collagens are known to be long-lived [1], it is possible that collagen VI may have accrued structural damage (e.g. from ROS, proteases or glycation) over time which may be reflected in the more globular regions of its microfibrillar ultrastructure. Collagen VI is particularly sensitive to MMPs-2 and -9, of which MMP-2 in upregulated in aged and degenerated IVD [46, 47]. Alternatively, the beaded region of the collagen VI microfibril is known to interact with numerous matrix proteins such as the proteoglycans biglycan and decorin and matrilins-1, -3 and -4, which in turn link to fibril forming collagen II and aggrecan to form an extensive ECM network [44]. Differences in peptide yield could reflect a change in the interaction state of aged microfibrils compared to young, which may affect the availability of interactive regions in the bead to trypsin digestion with functional consequences to known cell-collagen VI communication in the OAF [48].

Versican is a highly interactive proteoglycan, which together with collagen VI, exists as an integral component of translaminar cross bridges in the AF [49]. Previous analysis of this dataset demonstrated a significantly lower relative abundance for versican in aged OAF compared to young [31]. We reveal that two specific 50 aa segments of versican yielded more peptides in young OAF than in aged across all tissue regions, located at the interface between the GAGα and GAGβ regions **(Fig. 3B)**. Versican exists as four different isoforms in tissues with the V0 isoform containing both GAGα and β regions and the V1 and V2 isoforms containing only GAGα and GAGβ regions respectively [50]. The distribution of these isoforms was previously characterised in foetal IVD, where V0 was the predominating isoform [51], and in aged adult IVD, where the V1 isoform was reported as most prevalent (although the data was not shown) [52]. The higher peptide yields located at the interface between these GAG regions in young may indicate changing distributions of these versican isoforms with age, which may have functional implications.

CILPs-1 and -2 are monomeric ECM glycoproteins which have been implicated both in osteoarthritis [53, 54] and IVD degeneration [55, 56]. Although CILP1 expression was shown to increase both in articular cartilage and IVD with age [57, 58], we show that structure-associated differences in CILPs-1 and -2 also exist within the AF **(Fig. 3C, D)**. Several segments along the structures of CILPs-1 and -2 displayed either significantly higher or lower peptide yields in aged compared to young. Due to these age-dependant fluctuations, which were markedly conserved between all tissue regions, this perhaps reflects a more global change in the higher order structure of these proteins than seen for COL6A3 and versican which may affect function.

The observation that these four exemplar matrix proteins **(Fig. 3)** all displayed remarkable consistency between posterior, lateral and anterior regions of the OAF in their peptide yield patterns is important, as it provides clear evidence of tissue-wide mechanisms and consequences of ECM ageing. Despite this, a wide variety of proteins (ECM and non-ECM) were identified with tissue region-specific significant differences in peptide yield across their structures **(Fig. 2)**. A subset of proteins even displayed peptide yield patterns that were tissue region-conserved in parts of their structures but unique in others **(Fig. 4)**. This is interesting as it may indicate a superimposition of both tissue-wide and region-specific mechanisms or consequences of ageing which are acting upon the same proteins and may be reflected in the peptide yield patterns. These proteins include the alpha-1 chain of the collagen VI (COL6A1), the large tetrameric protease inhibitor alpha-2 macroglobulin (A2M), the beta chain of the clot-forming fibrinogen (FGB) and the immune response-regulating complement C3.

**Figure 4.**
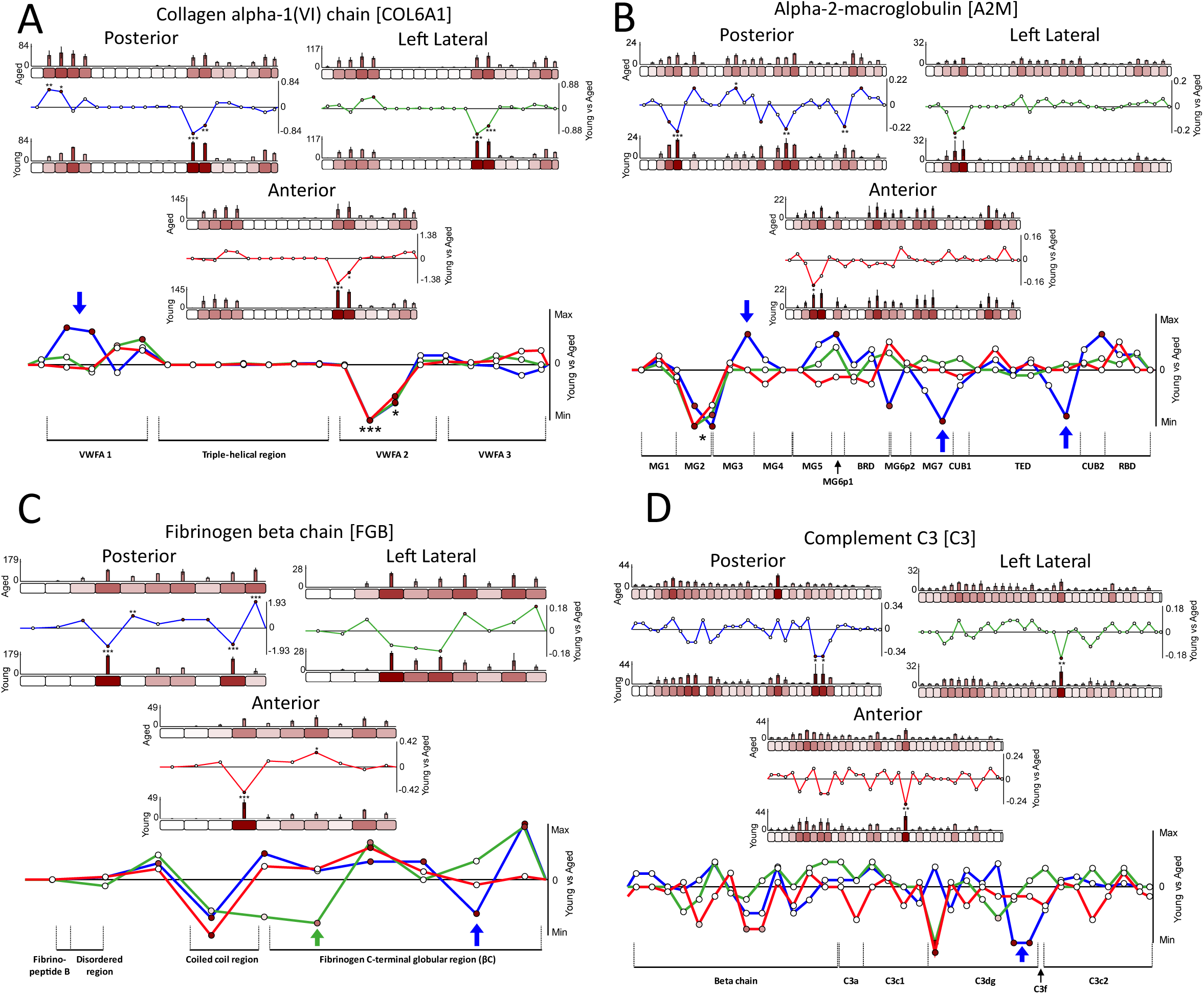
Proteins displaying both tissue region-conserved and -specific differences in peptide yield across their modular structures between young and aged IVD OAF. LC-MS/MS-detected peptide sequences were quantified within each 50 aa segment (bar graphs = average, normalised PSMs; error bars = SD). Differences in peptide yields across protein structures were assessed by subtracting average, normalised PSMs per segment in young from aged (Line graphs = aged-young PSMs/segment length; below zero line = higher in young, above zero line = higher in aged) and statistically compared (unpaired Bonferroni-corrected, repeated measures ANOVAs: *, p ≤ 0.05; **, p ≤ 0.01; ***, p ≤ 0.001; composite line graphs: stars = significant in all tissue regions; coloured arrows = tissue region-specific differences; aa ranges of Uniprot-sourced domains indicated). Two segments coinciding with the VWFA2 domain of COL6A1 **(A)** displayed higher peptide yields in young than in aged for all three OAF tissue regions. However, two segments near the N-terminal end of the protein (corresponding to VWFA1) exhibited significantly higher peptide yields in aged than in young that were posterior-specific. Similarly, an N-terminal protein region corresponding to the macroglobular (MG) 2 domain of A2M **(B)** displayed significantly higher peptide yields in young than in aged that were conserved between tissue regions. However, significant differences were also observed for three posterior-specific segments within the MG3 (higher in aged), MG7 and thioester (TED) domains (higher in young). FGB **(C)** had one segment within the coiled coil region of the protein which consistently displayed higher peptide yields in young than in aged for all three tissue regions, but also in one left lateral-specific segment (not significant) and one posterior-specific segment, located at either end of the globular (βC) region. Three segments within complement C3 **(D)** had significantly higher peptide yields in young than in aged OAF; one which was both anterior- and left lateral-specific, coinciding with the N-terminal half of the C3dg degradation product, and two which were posterior-specific, at C-terminal half of the same fragment.

The tissue-region conserved, significantly higher peptide yields observed in the VWFA2 domain of young COL6A1 compared to aged OAF **(Fig. 4A)** is analogous to that seen in the VWFA11 domain of COL6A3 **(Fig. 3A)**, both located just downstream of the triple helical region. Aged COL6A1 also exhibited a significantly higher peptide yield within the N-terminal VWFA1 domain compared to young, which was specific to the posterior region of the OAF **(Fig. 4A)**. As with COL6A3, both these globular regions of COL6A1 also correspond to the double beaded ultrastructural region of its constituent collagen VI microfibril. The observation that two out of three major alpha chains of collagen VI had structure-dependent differences closely associated to the microfibril bead further corroborates the hypothesis that damage to this long-lived ECM assembly may accrue with age or that the interaction state of aged microfibrils may differ compared to young. Although this appears to be predominantly tissue-wide (manifesting as conserved peptide patterns in all three tissue regions), posterior-specific mechanisms or consequences of ageing may also exist within the collagen VI microfibril.

Age-associated differences in protein structure, were not limited to ECM proteins as seen for A2M. This large, tetrameric glycoprotein is capable of non-specifically inhibiting all four classes of proteinases [59], including ECM-remodelling MMPs [60] and ADAMTSs [61, 62]. This inhibition is triggered via the cleavage of a “bait” region within A2M by the target protease which initiates a conformational change within the A2M complex, leading to the entrapment of the protease in a dimeric cage [63]. Tissue region-conserved, higher peptide yields in young OAF compared to aged were observed in the MG2 domain **(Fig. 4B)**, which is buried within the centre of the A2M monomer and attaches laterally to the CUB region [64]. Two posterior-specific regions, which coincide with the MG7 and TED domains of A2M, also yielded significantly more peptide in young than in aged OAF. These two domains, along with MG2, all frame an entrance to the central cavity of the dimer [64]. Since the tetrameric complex of A2M exists in two conformational states (protease bound or unbound), and since these domains frame the entrance to its entrapping cavity, the structure-associated differences observed here may reflect changes in these states between young and aged. This could represent a dynamic shift in global protease inhibition in the IVD because of ageing.

Fibrin fibres form the scaffold for blood clots, which comprise of activated fibrinogen dimers. Each monomer is composed of three N-terminal coiled chains (alpha, beta and gamma). Tissue region-conserved differences in peptide yields (higher in young than aged) were observed in the coiled coil region of the beta chain (FGB), whereas its globular region exhibited differences that were unique to the lateral and posterior regions of the OAF **(Fig. 4C)**. This C-terminal globular region (found in both beta and gamma chains [65]) is capable of multimeric association during fibrin protofibril formation [66]. Therefore, these structural differences between young and aged FGB, may reflect the distinct states of fibrin bundle formation within different regions of the IVD. Interestingly, A2M is able to inhibit the degradation of fibrin through its inhibition of plasmin and kallikrein [67]. As such, structure-associated changes in both A2M and FGB may indicate a potential for differing states of fibrin clotting in the aged IVD OAF compared to young.

Complement C3 has been described as the swiss-army knife of both innate and adaptive immunity. Its native form goes through a series of activation stages with each product and fragment released playing a unique functional role, including the opsonisation of foreign antigens, amplification of the complement response and immune cell recruitment [68]. PLF revealed significantly higher peptide yields within three segments of the C3 structure in young OAF compared to aged **(Fig. 4D)**, one which was both anterior- and lateral-specific and two unique to posterior. These segments all fall within the region of the protein’s C3dg degradation product, produced when C3b (the active form of C3) binds to complement receptors, to form C3c which is devoid of the C3dg region [69]. While C3c is released from cell, the C3dg fragment remains bounds and exerts a variety of effector functions including immune modulation [68]. The significant differences in peptide yields corresponding this fragment may reflect distinct activation states of complement C3 between young and aged OAF, potentially revealing clues to a changing immune response in ageing. The possibility that PLF could be used to assess the activation state of proteins in standard label-free LC-MS/MS datasets opens a potential new avenue for future study.

In addition to demonstrating differences in structure which were either conserved or unique to all three OAF regions tested, the specific peptide yield patterns displayed by these proteins may serve as fingerprints of particular mechanisms or consequences of ageing, as previously shown [27]. Crucially, this was not limited to ECM proteins alone, as these fingerprints were also used to investigate an extracellular protease inhibitor, coagulation protein and even a member of the complement pathway. Here, we demonstrate that patterns within these proteins, which coincided with functional regions, could provide clues as to their state within a macromolecular assembly (COL6A1, COL6A3, FGB), isoform composition (versican) or conformational (A2M) or activation (C3) state within OAF tissue, and how this may change with ageing.

### Collagens and basement membrane proteins among ECM components most affected by modifications to structure in ageing mouse lung

The mouse lung dataset was previously used for the development of an ageing lung atlas resource, comprised of both single cell transcriptomic and tissue proteomic analyses (https://theislab.github.io/LungAgingAtlas/) [30]. The proteomic analysis consisted of bulk whole lungs sourced from four young and four aged mice (3 and 24 months old). In addition to a global deregulation of protein expression with age, this formerly published study revealed extensive ECM remodelling between young and aged mice including alterations in the relative abundance of fibril-bridging (XIV, XVI), basement membrane (IV) and microfibrillar (VI) collagen alpha chains [30]. Although this dataset consisted of fractionated samples of soluble and insoluble proteins, we chose to focus on the ECM-rich fractions, which previously exhibited the largest separation of aged and young sample clusters by PCA [30] and are therefore most likely to harbour structure-associated evidence of ageing in long-lived matrix proteins.

MS/MS ion searches identified peptides corresponding to 668 proteins in both young and aged mouse lung groups (**Fig. S4 A**; peptide list: **Table S7**). PCA of peptides and their associated spectral counts demonstrated good separation of data between young and aged groups, with aged samples forming a more distinctive cluster compared to young samples **(Fig. S4 B)**. PLF analysis identified a total of 140 proteins displaying significant differences in peptide yields across their structures between aged and young groups (full PLF analysis results: **Table S8**). Classification analysis of these 140 shortlisted proteins revealed numerous collagens, elastic fibre-associated proteins and basement membrane laminins as the main ECM components structurally affected in ageing mouse lung **(Fig. 5)**. Architectural remodelling of ECM components in aged human lung has long been documented [34], including decreases in elastic fibre presence and an increased basement membrane thickness within alveolar walls. Here, we show for the first time that many proteins pertinent to these assemblies exhibit structure-associated modifications on a molecular level, providing clues as to specific targets of ageing mechanisms.

**Figure 5.**
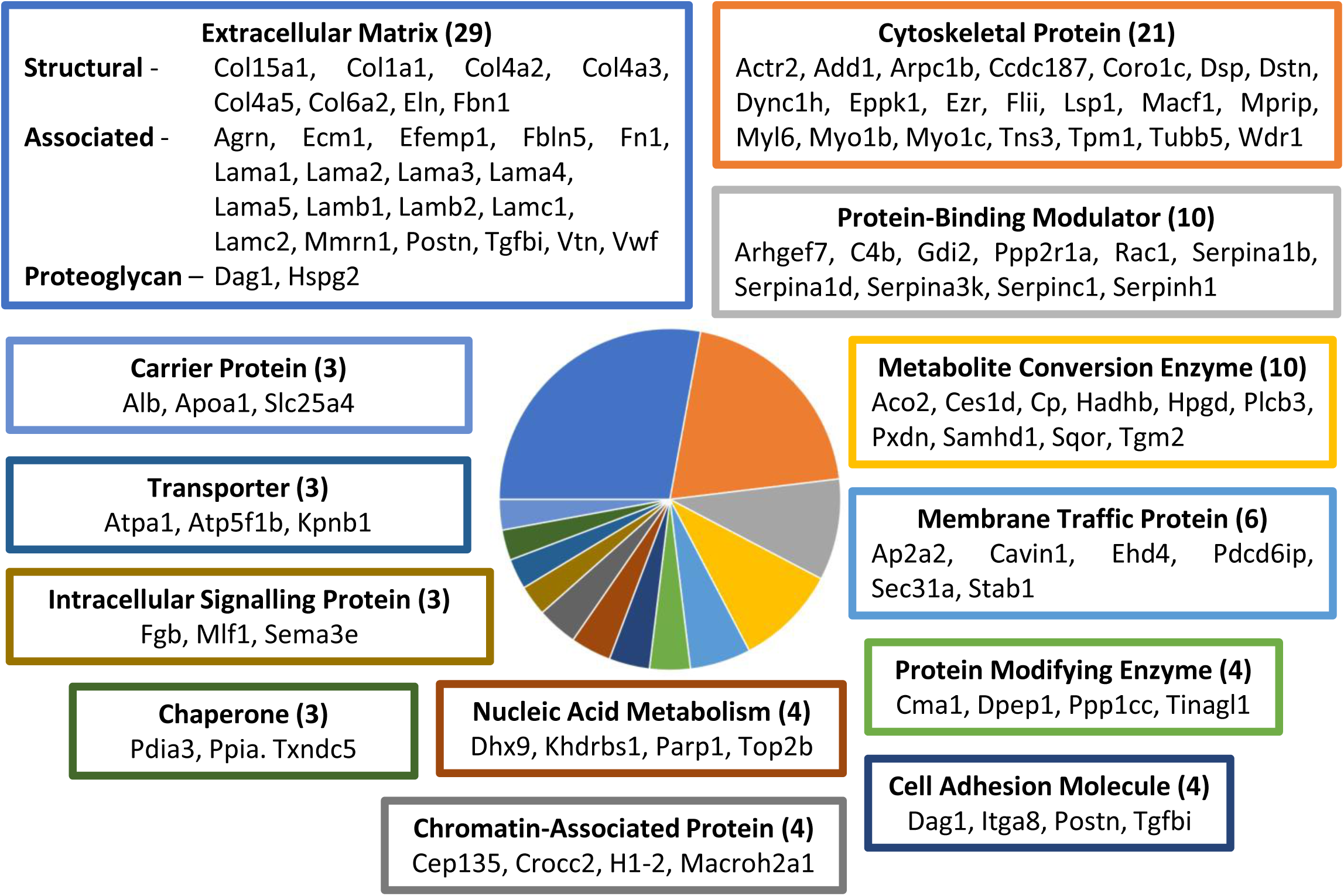
Classification analysis of proteins with structure-associated modifications reveals collagens and laminins as the main ECM components affected by ageing in mouse lung. PANTHER classification analysis indicated ECM proteins, cytoskeletal proteins (e.g. myosins), protein-binding modulators (e.g. serpins) and metabolite interconversion enzymes (e.g. transglutaminase 2 and matrix peroxidasin) as the top four classes structurally affected by age (classes and their protein identities are displayed with coloured frames matching their respective slice within the pie chart; minimum 3 proteins per class; total number of proteins per class indicated in brackets).

As performed for ageing human IVD **(Figs. 3, 4)**, specific age-dependent fluctuations in peptide yields across the structures of key lung ECM proteins were related to biological domains or regions in order to interrogate the functional consequences of their unique structural changes in ageing **(Fig. 6)**. These were the alpha 1 chain of the major structural collagen I (Col1a1) and four basement membrane component proteins including the alpha 2 chain collagen IV (Col4a2), laminin alpha chains -3 (Lama3) and -5 (Lama5) and heparan sulphate proteoglycan (HSPG) 2 core protein (perlecan).

**Figure 6.**
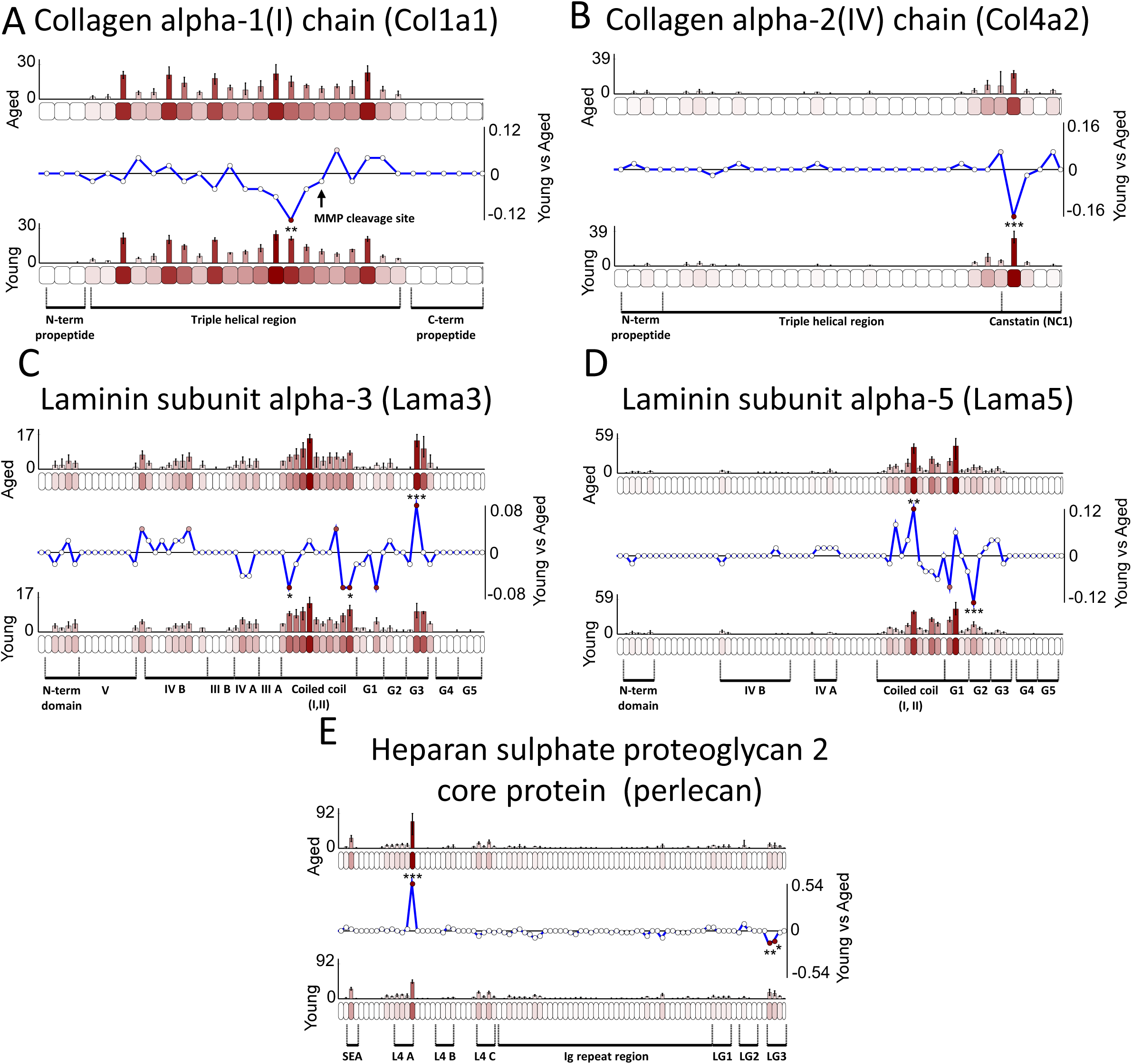
Key lung ECM proteins displaying significant differences in peptide yield across their modular structures between young and aged mice. LC-MS/MS-detected peptide sequences were quantified within each 50 aa segment (bar graphs = average, normalised PSMs; error bars = SD). Differences in peptide yields across protein structures were assessed by subtracting average, normalised PSMs per segment in young from aged (Line graphs = aged-young PSMs/segment length; below zero line = higher in young, above zero line = higher in aged) and statistically compared (unpaired Bonferroni-corrected, repeated measures ANOVAs: *, p ≤ 0.05; **, p ≤ 0.01; ***, p ≤ 0.001). Ranges of Uniprot-sourced regions are indicated. A single segment in Col1a1 **(A)** yielded significantly more peptides in young mouse lung than in aged, near the centre of the triple helical region of mature collagen I and upstream of a prominent MMP cleavage site. Similarly, Col4a2 **(B)** also contained a segment which yielded significantly more peptides in young near the C-terminal end of the protein, in a region corresponding to its canstatin cleavage product (NC1). Both Lama3 **(C)** and Lama5 **(D)** had significant differences in peptide yield within their coiled coil regions (higher in young for two Lama3 segments and higher in aged for one Lama5 segment). Lama3 and Lama5 also had segments within the globular (G) 3 and G2 domains which exhibited significantly higher and lower peptide yields in aged respectively. Two specific regions of perlecan **(E)** contained segments yielding significant differences in peptides: the laminin IV-like (L4) A domain (higher in aged) and the laminin G-like (LG) domain 3 (higher in young).

As the main, long-lived structural ECM scaffold component present in almost all connective tissues, the age-dependent remodelling of collagen I is well documented [70]. In healthy lung however, most ageing studies failed to identify changes to collagen presence (see reviews: [21]) and although the relative proportions of collagen I to III was shown to decrease in aged rats [71], this was not reproduced in mice [72]. In support of this, previous quantification analysis of the dataset analysed here found that the abundances of most lung collagens I and III did not change between aged and young mice [30, 73]. Despite this, we identified a segment within Col1a1 displaying a significantly higher peptide yield in young than in aged **(Fig. 6A)**, indicating that PLF may provide a more sensitive means of characterising age-associated changes to collagen I, which may occur on a more molecular level. This difference was seen in the triple helical domain of mature collagen I, suggesting that these modifications may be as a result of damage accumulation to structure. Levels of MMPs were previously shown not to change in this dataset [30], and activity of MMP-2 and -9 were shown to decrease with age in rat lungs [74]. Hence, this suggests that the structure-associated differences observed here for Co1a1 in mouse lung may not be due to changes in the expressions of secreted MMPs with age, in contrast to IVD. In support of this, the differences in Col1a1 peptide yield observed did not appear to coincide with its most prominent MMP cleavage site (**Fig. 6A**, black arrow). The presence of AGEs and related crosslinks in rat lung collagen however was previously shown to increase with age [75], with decreases in collagen solubility reported as a result [76]. As solubility is thought to play central role in the regional digestibility of ECM proteins [26, 29], the differences observed in the peptide yield patterns of the Col1a1 structure may instead be as a result of AGE accumulation with ageing.

Previous quantification analysis of this dataset showed higher levels of Col4a3 aged mice compared to young [30], indicating potential changes to collagen IV presence in ageing lung. Here we reveal that Col4a2 yielded significantly more peptides in young than in aged **(Fig. 6B)** within in a protein region corresponding to canstatin (NC1). Interestingly, this domain is released as a functional fragment (matrikine) during the proteolytic degradation of collagen IV by membrane-anchored (MT) MMPs-14 and -15 [77, 78]. Although previous analysis showed that no change in the levels of secreted MMPs [30, 73], the MT-MMPs [79] were not identified. It is possible that changes in MMPs 14 and 15 exist because of ageing which may lead to differences in the levels of this fragment, reflected here. Canstatin has been shown to actively inhibit cell proliferation, apoptosis and angiogenesis is therefore presumed to play a role in tissue homeostasis, with important implications for cancer therapy [78]. As such, decreasing levels of this fragment with age may be an important driver of ageing and age-related diseases in lung.

Levels of Lama3 and Lama4 were recently shown to be significant decreased in decellularized aged mouse lung compared to young, including a reduction in the expression of both laminin proteins for resident cells which were subsequently cultured on these aged scaffolds [80]. Although previous analysis indicated no significant changes in laminin chain abundance in this lung dataset [30], PLF analysis revealed multiple segments within the coiled-coil region and G domains of Lama3 and Lama5 which exhibited significant differences in peptide yield between aged and young mice **(Fig. 6 C,D)**. The laminins exist as trimeric complexes of alpha, beta and gamma chains. These chains wrap around each other in their coiled-coil regions, which then split into three N-terminal finger-like structures capable of interacting with other matrix components. At the C-terminal base of this complex is a region made of five globular (G) domains formed by the alpha chain alone which is capable of binding cell integrins and syndecans [81]. Significant differences in the availability of epitopes within the G domains in particular could be indicative of impaired cell-mediated interactions as a consequence of ageing. Alternatively, several MMPs are capable of cleaving laminin (including the calstatin-producing MMP14 discussed earlier) and some have even been shown to release functional matrikines [82]. As with collagen IV, differences in the structures of Lama3 and Lama5 could also be as a result of changes in MT-MMP activity with age.

To our knowledge, changes in perlecan have yet to be reported in healthy lung ageing however, due to its interactions with a multitude of ECM components, cell receptors and even growth factors, this proteoglycan has been implicated in the orchestration of age-related fibrosis in various connective tissues [83]. These many interactions are protein region-specific, with different domains of perlecan exerting unique functional roles. Significant differences in the peptide yield along the structure of this protein were observed within two domains in particular: L4A (higher in aged) and LG3 (higher in young) **(Fig. 6E)**. The L4-containing region is capable of binding growth factors (e.g. PDGF [84] and FGF7 [85]) to modulate signalling in tissue, whereas the LG-containing region is a more prolific binder of ECM components (e.g. nidogens, fibulins, elastin, collagen VI, matriglycan [86]). Differences in digestibility of domains within these regions between young and aged mice could reflect a change in these interactions because of ageing. Interestingly the LG3 domain can be cleaved and released by BMP1 or cathepsin L, as a fragment known as endorepellin which is capable of modulating cell signalling [87, 88]. Significantly more peptides were observed at the perlecan LG3 domain of young lung compared to aged, potentially corresponding to an increased presence of this fragment. Since this matrikine has been implicated as a mediator of fibrosis, age-associated changes in its presence may be a key driver of disease phenotypes in the ageing lung.

Basement membranes are composed of a meshwork of networked collagen IV, laminins and perlecan [89] all of which displayed structure-associated differences as a consequence of ageing **(Fig. 6)**, providing further evidence of potential disruption at the alveolar barrier, the crucial interface between capillaries and alveoli. In addition to potential disruptions in protein interactions (e.g. perlecan), the unique peptide fingerprints displayed by these proteins may enable the identification of specific mechanisms or consequences of ageing, such as changes in specific protease activity (such as MT-MMP cleavage of collagen IV and laminins) and accumulations of AGEs (on collagen I).

### Conserved differences within protein regions observed between human IVD and mouse lung ageing

Despite drastically shorter lifespans (∼2.5 vs. ∼70 years), mouse models are commonly used to characterise progression and treatment of human ageing and disease [18]. A recent study showed that aged mice exhibit typical features of human IVD ageing and degeneration, including loss of disc height and bulging alongside molecular changes and behaviours linked to increased pain compared to young mice [90]. Similarly, another study found that alterations in lung function and micromechanics in ageing mice were similar to that in humans [91]. Despite this, due to the longevity of ECM components which dominate tissues like lung and IVD, it is unclear whether mice can successfully model structure-associated changes to matrix proteins in humans, over many years. To examine whether certain mechanisms or consequences of ageing are shared between mouse and human proteins, we compared those identified with structure-associated alterations in human IVD ageing with those in identified mouse lung.

Out of a total of 284 age-affected proteins in human IVD OAF (across posterior, lateral and anterior regions) and 140 proteins in mouse lung, 37 were shared between both species **(Fig. S5)**. Classification analysis of these shared potential biomarkers of both human and mouse ageing revealed ECM proteins, mainly collagens and laminins, as the class predominantly affected in both tissues, accounting for ∼40% of proteins shared between species **(Fig. 7)**. Interestingly, this reflects an age-associated remodelling of the ECM which exists within the connective tissues of both human and mouse, regardless of differences in lifespan.

**Figure 7.**
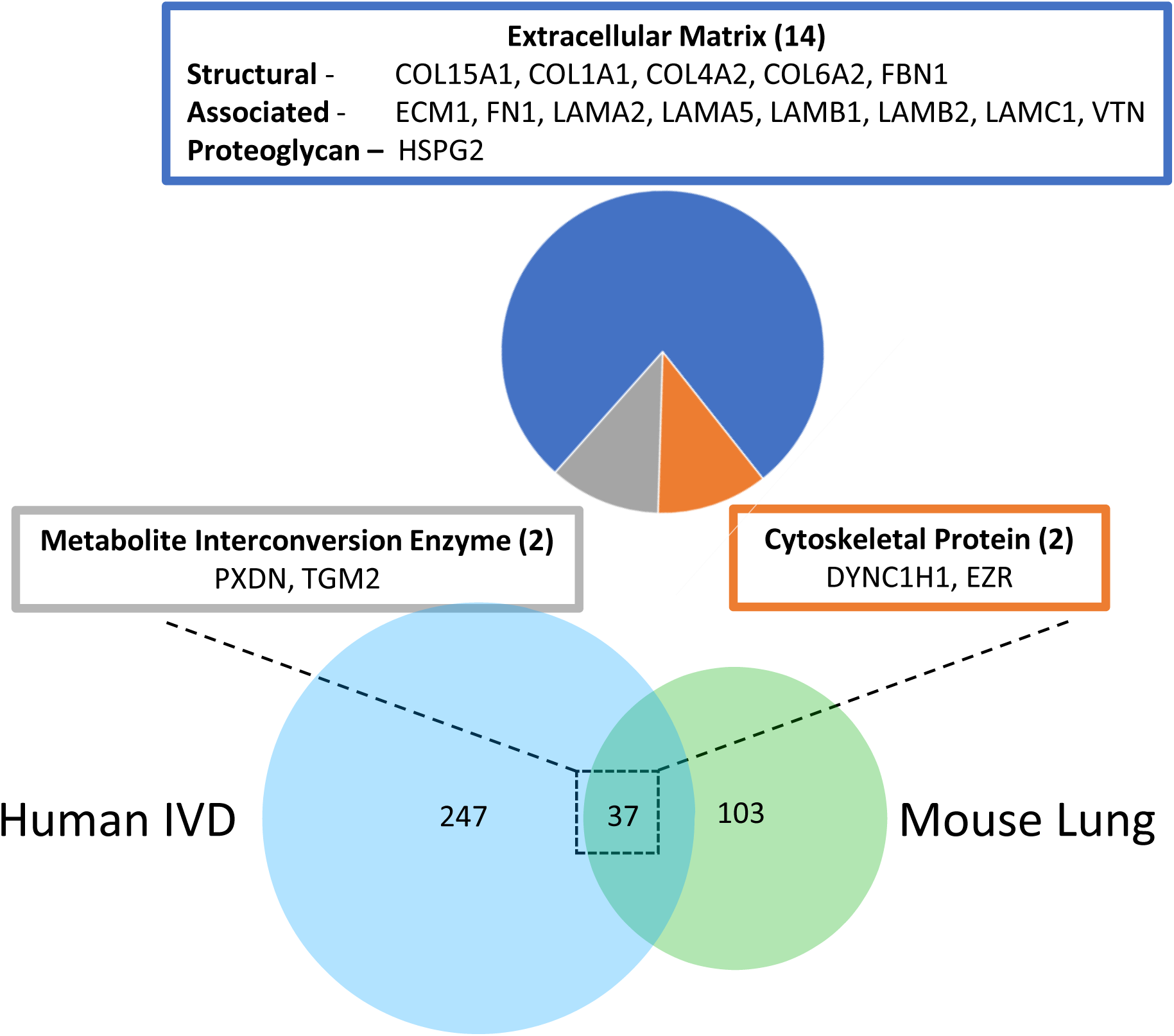
Classification analysis of common proteins in mouse lung and human IVD, identified with structure-associated modifications as a consequence of ageing. PANTHER classification of the 37 shared proteins identified ECM proteins (e.g. collagen and laminin chains, fibronectin and fibrillin-1), cytoskeletal proteins (dynein and ezrin) and metabolite interconversion enzymes (peroxidasin and transglutaminase 2) as the main classes structurally affected by ageing in both human IVD and mouse lung (classes and their protein identities are displayed; minimum 2 proteins per class; total number of proteins per class indicated in brackets).

Although these 37 shared proteins were identified as biomarker candidates of ageing in both human IVD and mouse lung, the mechanisms or consequences leading to those structure-associated changes may be unique to each. To determine which proteins may have evidence of the same ageing mechanisms or consequences that are irrespective of species and organs, peptide patterns across structures were compared to reveal protein regions of coinciding peptide yield differences. Of these 37 proteins, only four had regions exhibiting the same age-dependent differences in peptide yields for both human IVD and mouse lung: the adhesive ECM protein fibronectin, the basement membrane laminins beta-1 (Lamb1) and -2 (Lamb2) and the actin crosslinker, filamin-A **(Fig. 8)**.

**Figure 8.**
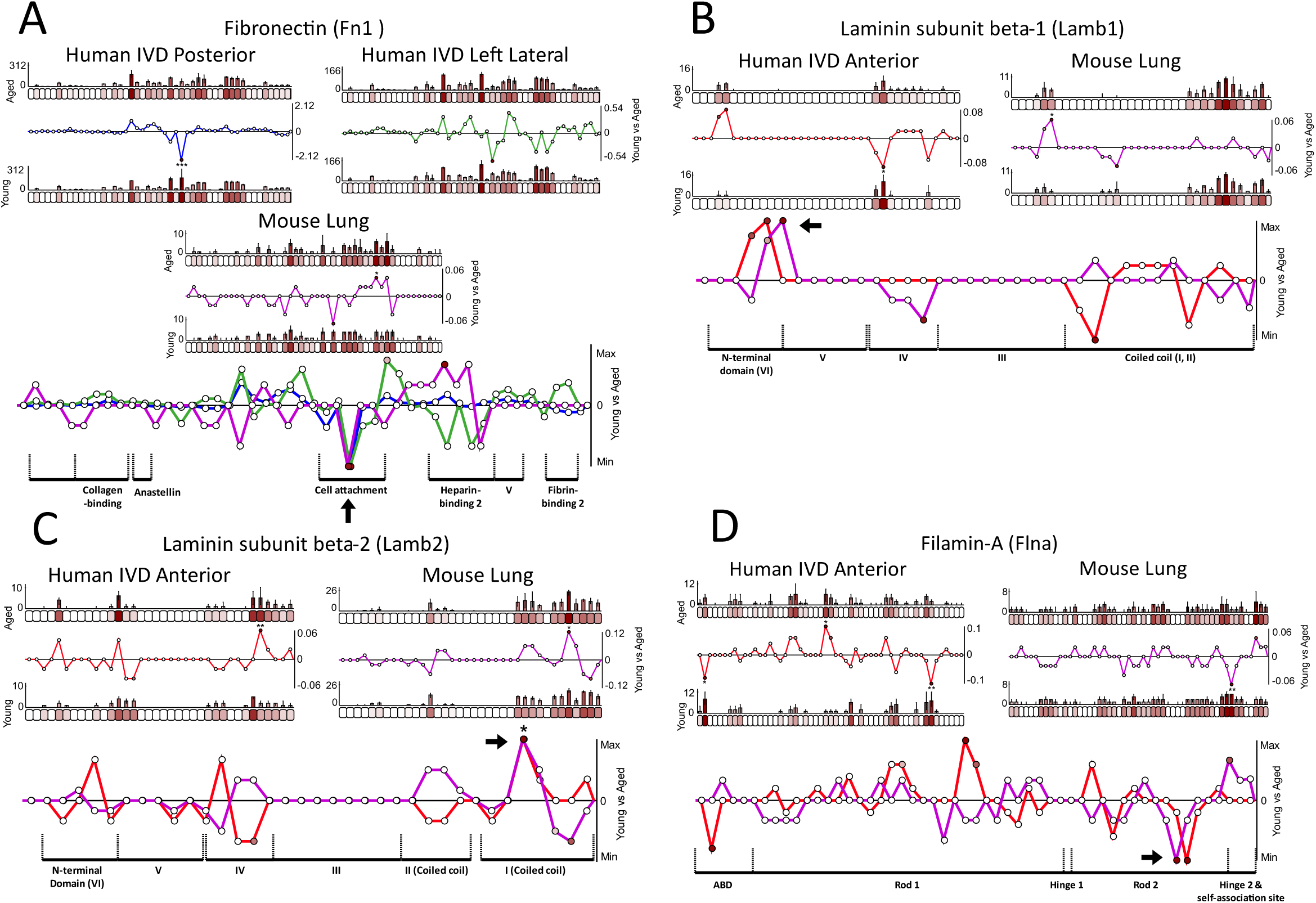
Proteins displaying species-conserved differences in peptide yield within regions of their modular structures between young and aged human IVD and mouse lung. LC-MS/MS-detected peptide sequences were quantified within each 50 aa segment (bar graphs = average, normalised PSMs; error bars = SD). Differences in peptide yields across protein structures were assessed by subtracting average, normalised PSMs per segment in young from aged (Line graphs = aged-young PSMs/segment length; below zero line = higher in young, above zero line = higher in aged) and statistically compared (unpaired Bonferroni-corrected, repeated measures ANOVAs: *, p ≤ 0.05; **, p ≤ 0.01; ***, p ≤ 0.001; composite line graphs: stars = significant in all comparison; black arrows = regions displaying species-conserved differences; aa ranges of Uniprot-sourced domains indicated). The same segment, near the centre of the fibronectin structure **(A),** exhibited similar higher peptide yields in young compared to aged human IVD (posterior and left lateral regions) as seen in young compared to aged mouse lung. This segment lies in the centre of a cell attachment region. Similar higher peptide yields for both aged human anterior IVD and aged mouse lung compared to young, were observed within the same N-terminal domain (VI) of Lamb1 **(B)**, and within the same C-terminal coiled coil domain (I) of Lamb2 (**C**). Filamin-A **(D)** also contained a region within its second rod domain which exhibited significantly higher peptide yields in both young mouse lung and human anterior IVD when compared to aged.

Fibronectin is a fibril-forming ECM glycoprotein, capable of a variety of cell and matrix interaction, which regulate cell adhesion, proliferation and activity [92] and elastic fibre assembly [93]. Aged mouse lungs demonstrate a decreased expression of fibronectin compared to young [80] and the ability of fibroblasts to secrete fibronectin was also shown to wane with age [94]. In contrast, higher levels of fibronectin were previously observed in aged bovine nucleus pulposus compared to young [35] and fragments of fibronectin have been detected in AFs from moderately degenerated IVDs [95]. The soluble form of fibronectin exists as a compact dimer connected at the C-terminus by a disulphide bond. Elongation of the dimer into fibrils is induced by integrin-binding of two central regions of the monomers after which clustering of fibronectin-bound integrins promotes the crosslinking of these fibrils into a stable matrix. Crucially, the significantly lower peptide yields seen consistently within a central region of fibronectin in both aged mouse lung and human IVD, compared to young, coincides specifically with this cell attachment region **(Fig. 8A)**. This indicates that connective tissue ageing both in humans and mice may have consequences to either fibronectin fibrillogenesis or general cell adhesion. Since these cell interactions are crucial to connective tissue homeostasis, these regional changes to fibronectin may play a prevalent role in its age-dependent disruption regardless of species.

In addition to two laminin alpha chains **(Fig. 6 C, D)**, structure-associated differences were also identified in two beta chain **(Fig. 6 B, C)** from aged mouse lung when compared to young, further evidencing a wider disruption of this basement membrane assembly. In addition, these higher peptide yields, within the N-terminal domain of Lamb1 **(Fig. 6B)** and within the coiled coil (I) domain of Lamb2 **(Fig. 6C),** were consistently seen in both aged human anterior IVD and aged mouse lung when compared to young. Although the role of laminins as basement membrane components is well characterised in lung [89], their function in the AF of the IVD is more elusive. They are widely present in the pericellular matrix of the IVD nucleus pulposus, however their distribution in the AF is more limited, especially in adult tissues [96]. Regardless of the unique roles these complexes may play in each tissue, the same protein regions were demonstrably affected in both species, potentially indicating a conserved mechanism or consequence of ageing. The N-terminal domain is critical for the self-assembly of laminins, their interactions with other ECM assemblies and their incorporation into basement membranes [97]. It is possible that the differences seen within the structures of Lamb1 and Lamb2 reflect a shift in their interaction states with similar consequences to both tissues.

Filamin-A is an important crosslinker of orthogonal actin fibrils (F-actin), involved in the dynamic remodelling of the actin cytoskeleton which governs cell motility, locomotion and mechanical resistance [98]. Filamin-A exists as an L-shaped homodimer (linked at the C-terminal end) and contains two actin-binding regions, one near the N-terminus (ABD) another further along the protein within the rod-1 domain [99]. As such, the dimer binds two F-actins at a 90° angle with its rod-1 domains resting flat along the filaments [98]. Similar regions within the rod 2 domain of filamin-A yielded significantly more peptides in young than in aged, for both mouse lung and human anterior IVD **(Fig. 8D)**. Interestingly, this domain functions to bind integrins, linking the actin cytoskeleton to the cell membrane and in turn to the ECM matrix [100]. As such, differences within this region may reflect a changed state of resident cell interactions with the ECM, which remains consistent between species and organs.

Interestingly, as components of the pericellular matrix, fibronectins and laminins are both integral to cell adhesion while filamin-A plays a role in anchoring the actin cytoskeleton to the outer ECM [100]. This suggests perhaps that the conserved structure-associated differences, seen in both mouse lung and human IVD, are connected to a changing cellular microenvironment during connective tissue ageing. To our knowledge, this study is the first to provide evidence that ECM proteins are subjected to similar structure-associated differences in mouse and human ageing, irrespective of their vastly different lifespans and tissue functions. It indicates that universal markers of connective tissue ageing may exist within the ECM and provides clues into mechanisms or consequences which may be conserved and therefore crucial to our understanding of ageing.

### Sex-conserved and -unique differences identified within protein structures from human atherosclerotic plaque

The human atherosclerotic dataset was previously used for the proteomic analysis of distinct arterial lesions and includes label-free LC-MS/MS of plaque core, fatty streak and internal control tissue sourced from ten aged males and females [32, 33]. Former study of this dataset revealed sex-conserved differences in the relative abundance of ECM components, such as proteoglycans mimecan and biglycan, which are indicative of ECM remodelling in atherosclerosis [32]. For this study, we chose to investigate plaque core tissue specifically (in comparison to internal control) as it exhibited the largest differences in proteomic composition [32], and therefore disease progression. Since sex-specific differences in abundance were observed previously for this dataset in key players of disease progression (e.g. fibrinogens and apolipoproteins) [32, 33], we too stratified this comparison to males and females in order to further investigate similarities and differences between sexes.

Peptides corresponding to 510 proteins in males and 461 proteins in females were identified in both atherosclerotic plaque and control artery by MS/MS peptide ion searches (**Fig. S6**; peptide lists: **Tables S9, S10**). PCA of peptides and their associated spectral counts revealed good separation of data between plaque and control artery for both male and female groups **(Fig. S7)**. Male plaque samples in particular formed a distinct cluster in comparison to other groups, suggesting sex-specific homogeneity. PLF analysis led to the identification of 258 proteins across both sexes exhibiting significant differences in peptide yields across structures between plaque and control artery, 64 of which were shared between males and females **(Fig. S8)**.

Classification analysis of these shortlisted proteins once again identified ECM proteins as the major class structurally modified in arterial atherosclerosis, comprising ∼20% of affected proteins shared between male and females **(Fig. 9)**. This included chains from two basement membrane collagens IV and XVIII, the fibrillar collagen-bridging collagen XII and the microfibrillar collagen VI as well as tenascins-X and -C and proteoglycans perlecan and mimecan. Previous label free quantification of this dataset identified biglycan and mimecan as the only ECM components with significantly altered relative abundances in plaque compared to control [32]. In contrast, PLF identified 35 affected ECM proteins across males and females, indicating a broader remodelling of ECM than initially observed, which was better reflected through structure-associated measurements.

**Figure 9.**
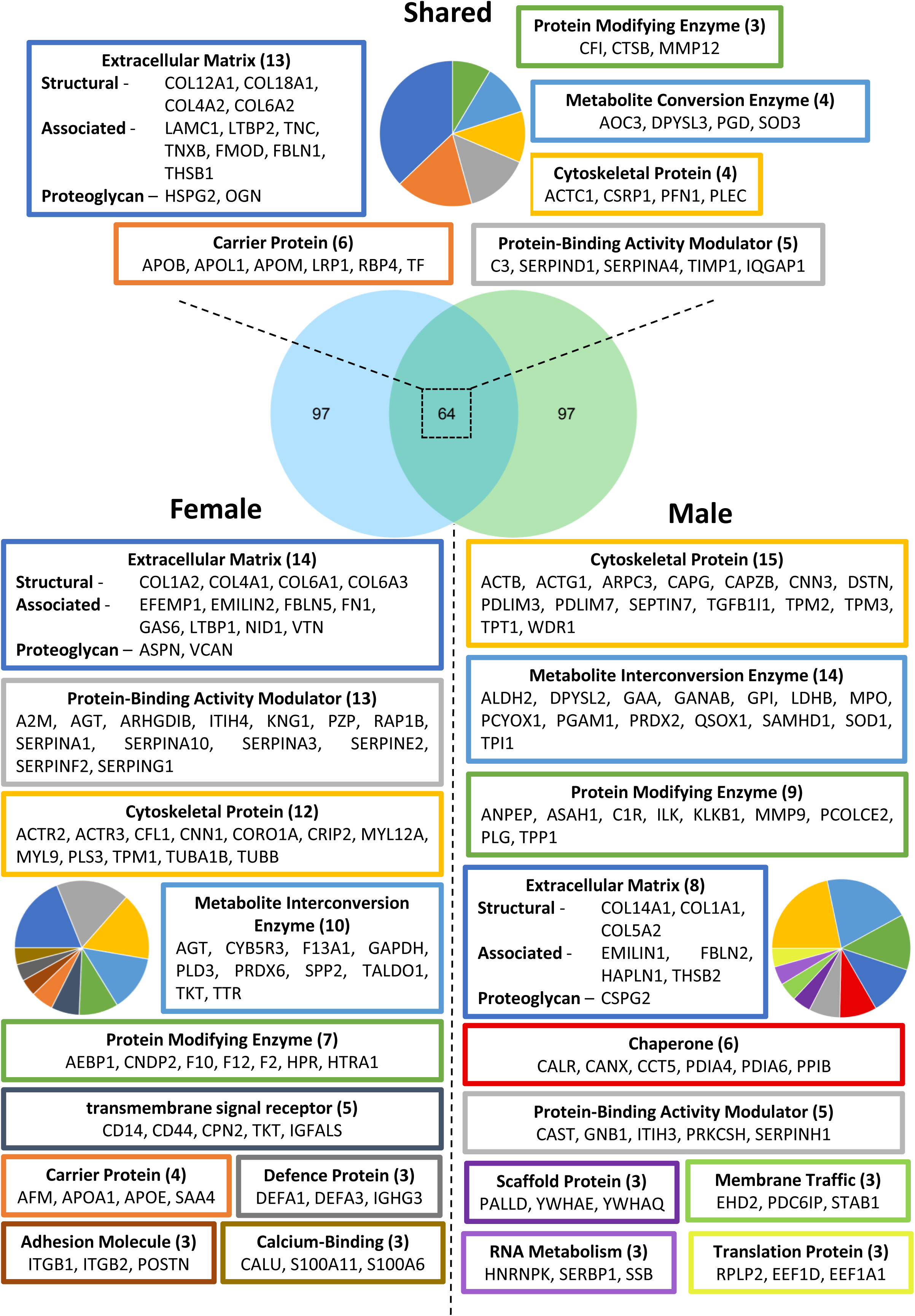
Classification analysis of proteins with structure-associated differences reveals ECM proteins, cytoskeletal proteins, protease and protease inhibitors and metabolic enzymes as the main classes affected in atherosclerosis. PLF analysis identified 161 proteins in males and in females with significant differences in peptide yield across structure (32% and 35% of protein identities, respectively; full PLF analysis results: **Tables S11, S12**.) PANTHER classification analysis [42] indicated ECM proteins as the main class affected in females and also between male and female groups (classes and their protein identities are displayed with coloured frames matching their respective slice within the pie charts; minimum 3 proteins per class; total number of proteins per class indicated in brackets). Cytoskeletal proteins were also one of the classes most affected across both sexes (top for male; e.g. actins and myosins), as well as metabolite interconversion enzymes (e.g. superoxide dismutases), protein modifying enzymes (e.g. MMPs and cathepsin B), and protease inhibitors (e.g. serpins and tissue inhibitor of metalloproteinase [TIMP]-1).

Peptide yield differences across the structures of PLF-identified proteins were further compared between males and females to investigate whether fluctuations between plaque and control arteries were sex-dependent. Additionally, significant differences within key proteins were related to biological domains or regions to reveal potential functional consequences to arterial atherosclerosis. Several key proteins had regions which yielded significantly more peptides in plaque than in control consistently between males and females. This included proteoglycans perlecan and mimecan, the cysteine protease cathepsin B and the actin crosslinking alpha actinin-4 (ACTN4) **(Fig. 10)**.

**Figure 10.**
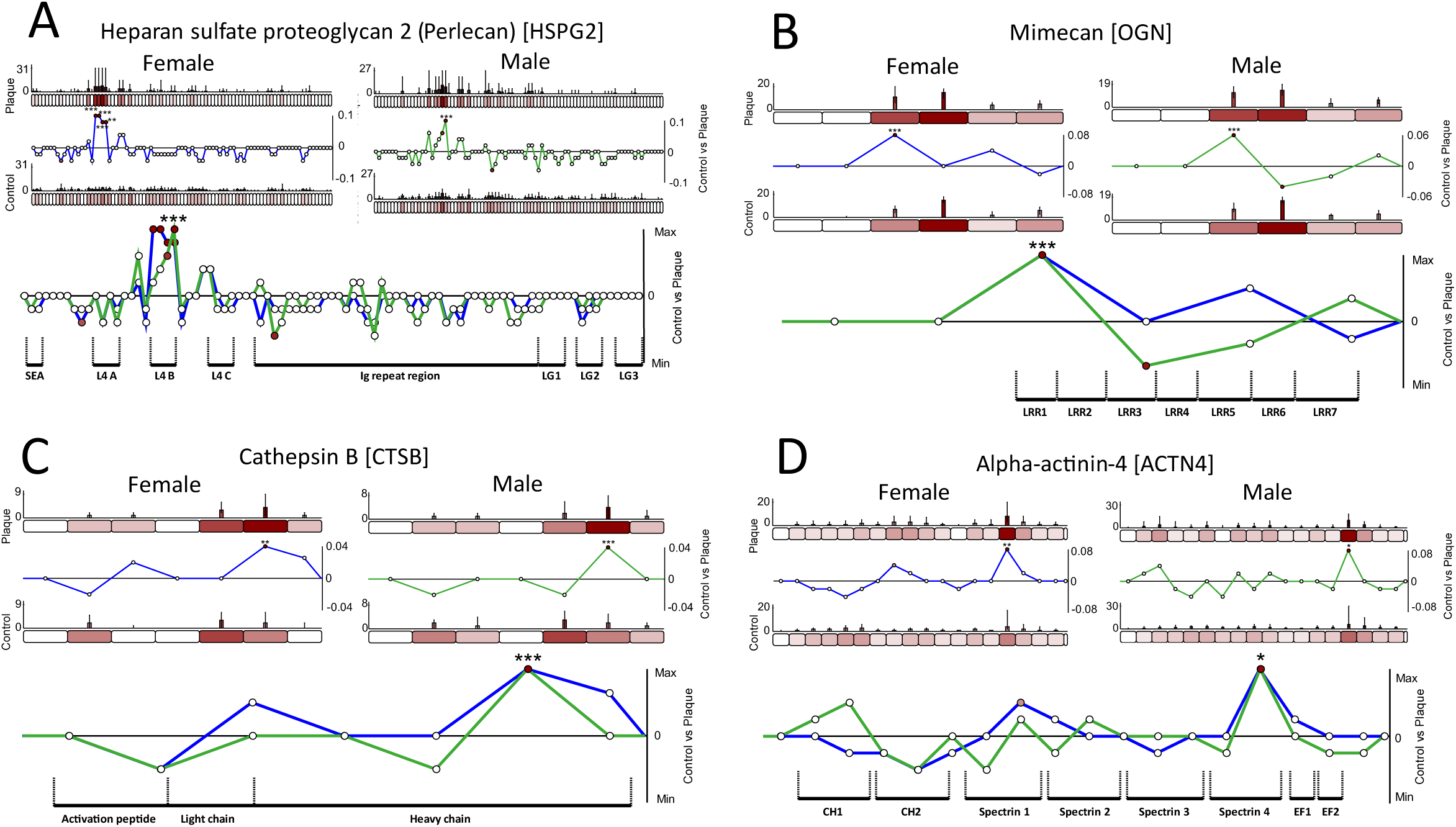
Proteins displaying sex-conserved differences in peptide yield across their modular structures between human atherosclerotic plaque and internal control artery. LC-MS/MS-detected peptide sequences were quantified within each 50 aa segment (bar graphs = average, normalised PSMs; error bars = SD). Differences in peptide yields across protein structures were assessed by subtracting average, normalised PSMs per segment in control from plaque groups (Line graphs = plaque-control PSMs/segment length; below zero line = higher in control, above zero line = higher in plaque) and statistically compared (paired Bonferroni-corrected, repeated measures ANOVAs: *, p ≤ 0.05; **, p ≤ 0.01; ***, p ≤ 0.001; composite line graphs: stars = significant in both females and males). Ranges of Uniprot-sourced regions are indicated. One segment within perlecan **(A)** and another within mimecan **(B)** displayed significantly higher peptide yields in plaque than in control that were consistent between females and males. These corresponded to the L4 B domain of perlecan and the first leucine-rich repeat (LRR) of mimecan. Cathepsin B **(C)** also contained a single segment near the C-terminal end of its heavy chain which yielded significantly more peptides in plaque than in control consistently between males and females. ACTN4 **(D)** had one segment within its fourth spectrin repeat, near the C-terminal end of the protein which also yielded more peptides in plaque than in control for both male and female groups.

HSPGs are capable of reducing the subendothelial retention of low-density lipoproteins (LDLs) within arteries by 1.5 - 10 fold [101]. As such, the reduction of perlecan on arterial walls is thought to contribute to the disease progression of atherosclerosis [102]. Here, we show that these changes are not limited to differences in protein abundance but may also be structure-associated. The higher peptide yields observed within both male and female plaque-sourced perlecan compared to control were very specific to the L4 B domain of the proteoglycan **(Fig. 10A)**. This pattern is similar to that seen for mouse lung ageing, where higher peptide yields were also seen within the L4 A domain of perlecan in aged compared to young **(Fig. 6E)**. This potentially indicates that the L4-containing regions of this protein may be particularly sensitive to structure-associated changes. This region interacts with the basement membrane ECM protein WARP and collagen VI [86] and is capable of sequestering growth factors [84, 85], as mentioned previously. Perhaps structure-associated differences within these L4 regions reflect disturbances in these interactions in both atherosclerosis and ageing, drawing parallels between the two processes.

Mimecan is a keratan sulphate proteoglycan that has been described as an emerging biomarker of atherosclerosis [103]. In human carotid plaques, its presence was recently shown to be downregulated in vascular smooth muscle cells but upregulated in certain plaque regions (close to collagen fibres and neutral lipids) [104]. The same study also showed a positive correlation between plaque levels of mimecan and age, with higher levels predicting future cardiovascular-related death. Previous analysis of mimecan within this proteomics dataset indicated significantly reduced levels in plaque core compared to control [32]. Here, we show that mimecan derived from both male and female plaque yielded consistently higher peptides within LRR1 when compared to control **(Fig. 10B)**. This structure-associated difference may reflect changes in mimecan activity and its protein interactions with potential consequences to collagen fibrillogenesis [105] which may provide clues to the mechanism of plaque formation.

Cathepsins B, X and D were previously shown to be upregulated in atherosclerotic lesions and play crucial roles in disease progression [106]. Part of this process is orchestrated by cathepsin B which is thought to progressively remodel and degrade the ECM in arterial intima through its secretion by resident macrophages [107]. The inactive form of the enzyme is localised within lysosomes, where it is proteolytically cleaved into active cathepsin B via removal of its activation peptide. It is then further processed into heavy and light chains, linked together by a disulphide bond. A higher peptide yield was observed near the C-terminal end of its heavy chain **(Fig. 10C)** for both male and female plaque compared to control. Since regional fluctuations in peptide yields were shown to correlate with differences in a protein’s quaternary structure [28]. Should this occur in an enzyme like cathepsin B, it may reflect differences in the state of its activity between plaque and control, via complexing with substrates or inhibitors. This would further corroborate the role of cathepsin B plays in the remodelling of ECM during plaque formation. The possibility that changes in the activity states of certain enzymes could be measured with PLF is an exciting prospect which requires further investigation and may provide an opportunity to gauge active proteolysis in addition to protein quantification.

Alpha-actinin exists as an antiparallel homodimer with its monomers associating along their spectrin repeats to form a cylindrical rod domain [108]. The N-terminal ends of this rod are flanked by two actin-binding CH1 and CH2 domains, enabling the formation of crosslinks between F-actin. As such, ACTN4 plays a crucial role in the regulation of the actin cytoskeleton, and therefore cell motility. Higher peptide yields were observed within the spectrin 4 domain of plaque-derived ACTN4 of both males and females, compared to control artery **(Fig. 10D)** indicating a potential change in this complex’s higher order structure. To our knowledge, the effect of atherosclerosis on ACTN4 has yet to be demonstrated, although exposure of oxidised LDLs (oxLDLs) to podocytes was recently shown to significantly enhance podocyte migration by increasing ACTN4 expression [109]. As a major crosslinker of actin, changes in ACTN4 are associated to alterations in cell motility and migration [110]. As such, structure-related differences within this protein may reflect an altered morphological state of resident plaque cells compared those in control artery.

Perlecan, mimecan, cathepsin B and ACTN4 all displayed regional significant differences in peptide yield, between plaque and control artery, that were consistent between males and females, providing clear evidence of sex-wide consequences of atherosclerosis **(Fig. 10)**. However, several proteins harboured structure-associated differences which were sex-specific **(Fig. 9)**, including some displaying peptide yield differences that were both consistent between male and females in some protein regions, but sex-specific in others. Interestingly, this may be indicative of distinct mechanisms or consequences acting upon different regions of the same proteins. This was observed in the ECM microfibrillar COL6A2 and the actin-crosslinking filamin-A **(Fig. 11)**.

**Figure 11.**
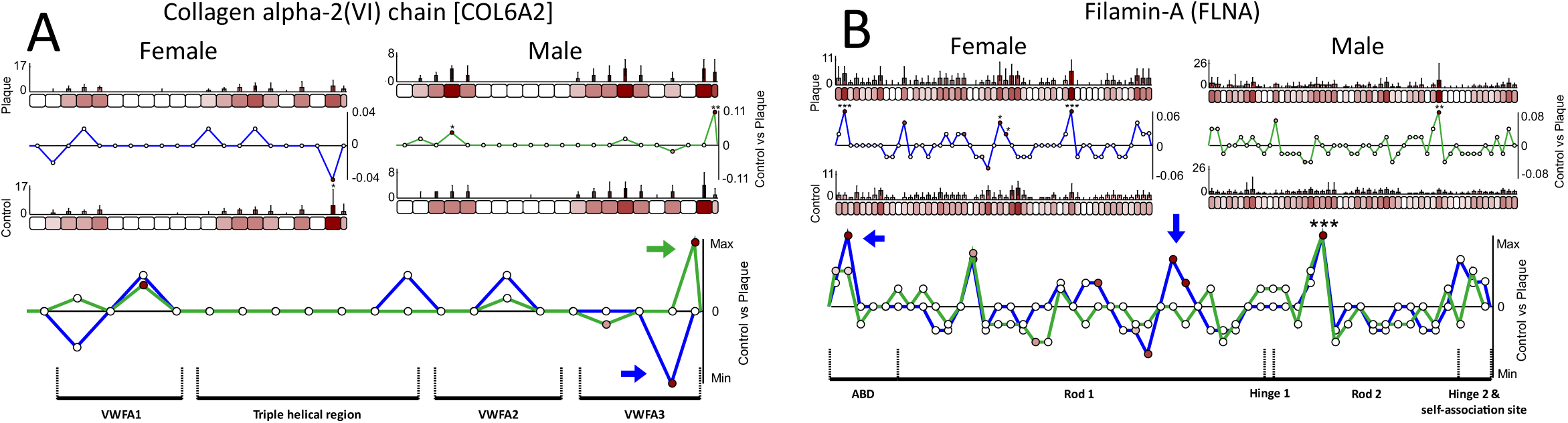
Proteins displaying both sex-conserved and sex-specific differences in peptide yield across their modular structures between human atherosclerotic plaque and internal control artery. LC-MS/MS-detected peptide sequences were quantified within each 50 aa segment (bar graphs = average, normalised PSMs; error bars = SD). Differences in peptide yields across protein structures were assessed by subtracting average, normalised PSMs per segment in control from plaque groups (Line graph = plaque-control PSMs/segment length; below zero line = higher in control, above zero line = higher in plaque) and statistically compared (paired Bonferroni-corrected, repeated measures ANOVAs: *, p ≤ 0.05; **, p ≤ 0.01; ***, p ≤ 0.001; composite line graphs: stars = significant in both females and males; coloured arrows = sex-specific differences; aa ranges of Uniprot-sourced domains are indicated). The fourth segment, located within the VWFA1 domain near the N-terminal end of COL6A2 **(A)**, displayed higher peptide yields in plaque than in control for both male (significant) and female groups. However, at the opposite C-terminal end, within the VWFA3 domain of the protein, its two final segments displayed significantly higher peptide yields in plaque for male but significantly lower yields for female when compared to control. Filamin-A **(B)** contained one segment within its second rod domain which yielded sex-consistent, significantly higher peptide yields in plaque than in control, but also one segment within its N-terminal actin binding domain (ABD) and two more segments in the C-terminal half of the first rod domain which displayed significantly higher peptides that were unique to plaques in females.

In artery and atherosclerotic lesions, collagen VI can be found interspersed between collagen I and III fibres in both the tunica media and subendothelium [111, 112]. Regional, significant differences in peptide yields between plaque and control were observed in protein segments located within the VWFA1 and 3 domains of COL6A2 **(Fig. 11A)**. As also observed for COL6A3 **(Fig. 3A)** and COL6A1 **(Fig. 4A)** in ageing IVD, changes within these VWFA domains correspond to the beaded regions of the collagen VI microfibril ultrastructure. This indicates that the highly interactive microfibril bead [44] may be particularly sensitive to mechanisms of change (or damage), as a consequence of either ageing of age-related disease. Although the direct involvement of collagen VI in the progression of atherosclerosis has yet to be determined, platelets are known to bind collagen VI indirectly through their association to the blood clotting-VWF protein (which in turn binds collagen VI) or directly to the platelet integrin α2β1 [112, 113]. Since platelet associations with the endothelium are thought to influence the progression of atherosclerosis [114], structure-associated changes within the collagen VI microfibril bead may have functional consequences to these interactions. As well as in ageing tissue, the progressive glycation of long-lived collagens, and the associated accumulation of AGEs, is also thought to influence the progression of atherosclerosis [115]. As mentioned, changes in the trypsin digestibility of regions within the microfibril bead may reflect an accumulation of AGEs.

Recently, filamin-A expression was shown to be elevated in human atherosclerotic plaques and, in the same study, its targeted reduction in macrophages led to their decreased migration and proliferation which ultimately reduced the size of atherosclerotic plaques in mouse models [116]. Interestingly, only female plaque-derived filamin-A exhibited higher peptide yields than control within the actin-binding domain (ABD) and rod 1 domain **(Fig. 11B)**, which may reflect a sex-specific difference in the actin cytoskeleton of cells in atherosclerotic lesions. The rod 2 domain however, displayed higher peptide yields in plaque that were consistent between both sexes. As mentioned previously, this domain binds to integrins [100], but also to platelet adhesion glycoprotein (GP) Ib-IX-V receptors [117] which links the actin cytoskeleton to the cell membrane. Changes within this domain may reflect an alternate state of the actin cytoskeleton within cells of atherosclerotic lesions, perhaps linked to migration and proliferation as observed previously in macrophages [116].

PLF analysis of plaque and control artery proteomes revealed that structure-associated changes are not limited to just ageing proteins, but also exist as a consequence of age-related diseases like atherosclerosis. These differences were not only observed in ECM proteins (perlecan, mimecan and collagen VI), but also cytoskeletal crosslinkers ACTN4 and filamin-A and in the ECM modifying protease cathepsin B, reflecting a global, semi sex-specific shift in matrix remodelling and cell adhesion.

### Conserved differences within protein regions observed between IVD ageing and arterial atherosclerosis

Atherosclerosis has been characterised as premature biological ageing, where mechanisms such as cellular senescence and its associated promotion of inflammation and perturbed maintenance of the ECM drive the progression of disease [19]. Links between atherosclerosis and IVD ageing and degeneration in particular have also been shown, with aortic plaque lesions, occluded lumbar arteries and high serum cholesterol levels all associated [118]. Concordantly, levels of oxidised (ox-) LDLs (accumulation of which in the vascular intima is a hallmark of atherosclerosis [119]) were also positive correlated with the progression degeneration in both the nucleus pulposus and OAF [120].

Despite these links, it remains unknown whether proteins are similarly affected by ageing in IVD and atherosclerosis. To examine whether certain mechanisms or consequences of ageing are shared in age-related diseases like atherosclerosis, we next compared the proteins identified with structure-associated alterations in IVD OAF ageing with those in identified in arterial atherosclerosis. Of the 258 proteins identified for atherosclerosis (in both male and female) and 284 proteins identified for IVD ageing (across posterior, lateral and anterior tissue regions) with structure-associated differences, 85 were shared **(Fig. S9)**. Classification analysis of these shared potential biomarkers **(Fig. 12)** identified ECM protein as the major class affected in both IVD ageing and arterial atherosclerosis, including multiple alpha chains of collagens (I,IV,V, VI, XIV), elastic fibre-associated proteins (EMILIN-1, fibulin-1, LTBP2), thrombospondins (−1, −2), tenascins (- C, -X) and proteoglycans (perlecan and versican). ECM proteins accounted for ∼24% of proteins identified with significant structure-associated differences in both ageing and age-related disease, indicating a shared matrisomal remodelling between IVD and artery.

**Figure 12.**
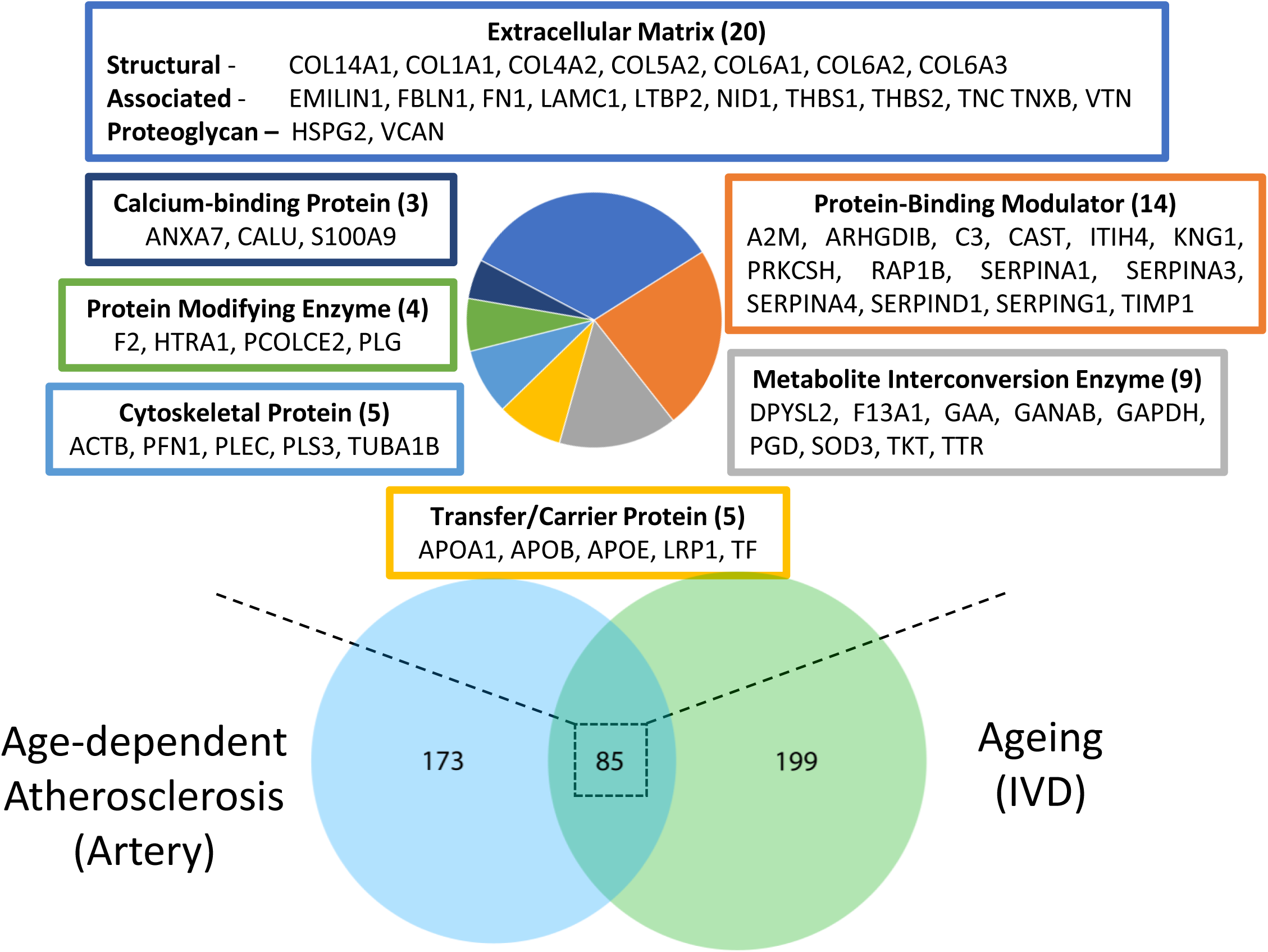
Classification analysis of proteins identified with significant structure-associated differences in both arterial atherosclerosis and IVD ageing. PANTHER classification analysis [42] identified several protein classes affected as a consequence of both age-related disease and ageing (classes and their protein identities are displayed with coloured frames matching their respective slice within the pie chart; minimum 3 proteins per class; total number of proteins per class indicated in brackets). As well as ECM proteins, this included protein-binding activity modulators (e.g. protease inhibitors A2M, serpins and TIMP1), metabolite interconversion enzymes (e.g. superoxide dismutase 3), transfer/carrier proteins (e.g. apolipoproteins) and cytoskeletal proteins (e.g. actin beta and tubulin alpha-1B) as the top five.

To determine whether any of these 85 common potential biomarkers had evidence of structure-associated mechanisms or consequences that may be shared between ageing and age-related atherosclerosis, peptide patterns across their structure were compared, revealing regions with coinciding peptide yield differences. Since elements of atherosclerosis have been viewed previously as “accelerated forms of vascular ageing” [19], we compared the peptide yield differences between aged and young IVD to the same differences between plaque and control. Four proteins were identified with regions yielding very similar peptide yield differences between aged vs. young IVD and plaque vs control artery: the protease inhibitor A2M, the coagulant prothrombin, the alpha-1 chain of the fibrillar collagen-bridging collagen XIV (COL14A1) and the LDL-associated apolipoprotein-B (APOB) **(Fig. 13)**.

**Figure 13.**
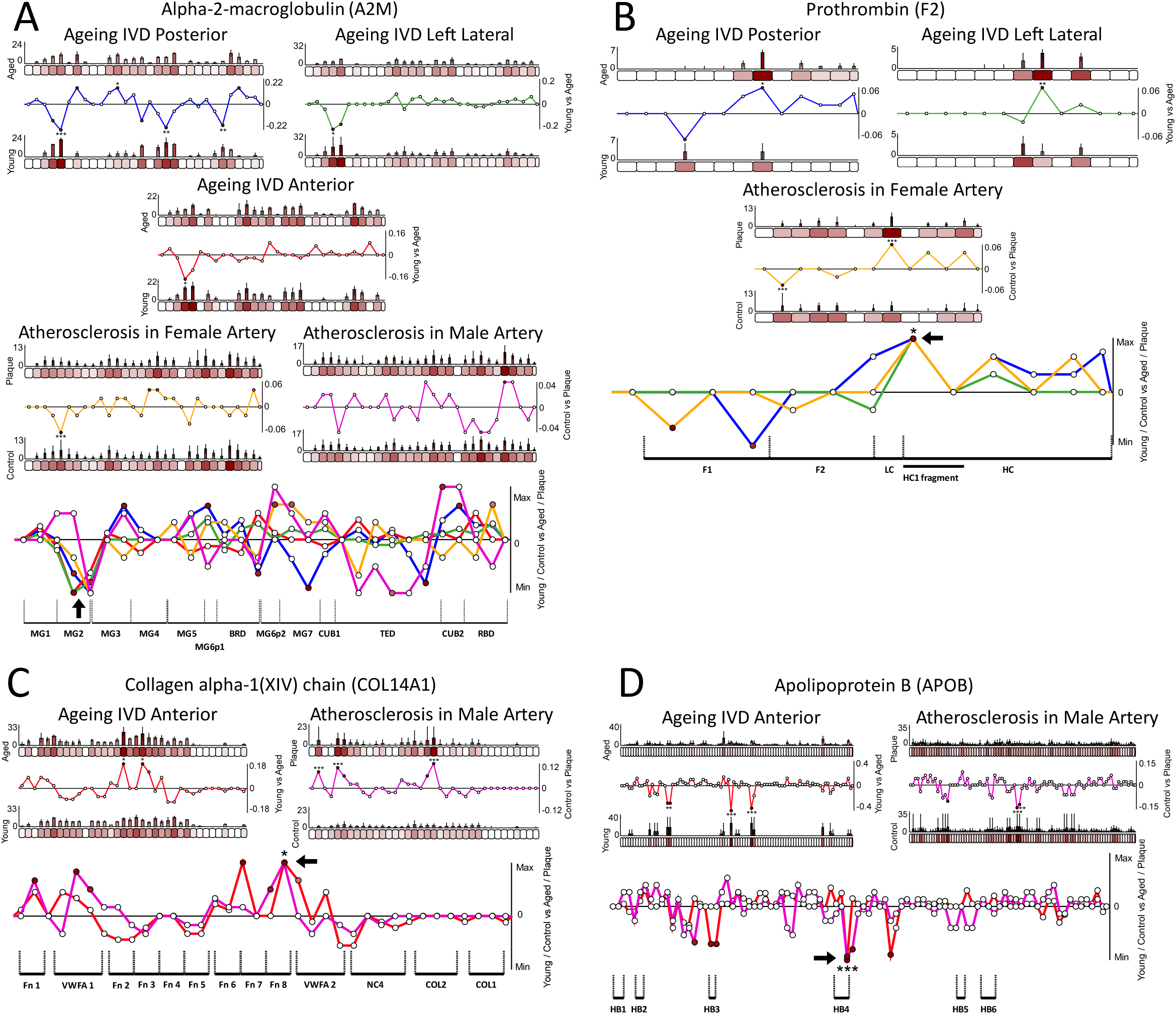
Proteins displaying conserved differences in peptide yields within regions of their modular structures between ageing in IVD OAF and age-related atherosclerosis in artery. LC-MS/MS-detected peptide sequences were quantified within each 50 aa segment (bar graphs = average, normalised PSMs; error bars = SD). Differences in peptide yields across protein structures were assessed by subtracting average, normalised PSMs per segment in young from aged for IVD and in control from plaque for artery (IVD line graph = aged-young PSMs/segment length; artery line graph = plaque-control PSMs/segment length; below zero line = higher in young/control, above zero line = higher in aged/plaque) and statistically compared (Bonferroni-corrected, repeated measures ANOVAs, unpaired for IVD, paired for artery: *, p ≤ 0.05; **, p ≤ 0.01; ***, p ≤ 0.001; composite line graphs: stars = significant in all comparison; black arrows = regions displaying ageing and atherosclerosis-conserved differences; aa ranges of Uniprot-sourced domains indicated). The MG2 domain of A2M **(A)** contained segments which exhibited the same higher yield of peptides in young than in aged, for all three tissue regions of the IVD (posterior, left lateral and anterior), and in control compared to plaque for both male and female arteries. These differences were significantly different for all comparisons except male artery. Prothrombin **(B)** had a single segment within its heavy chain (HC) which exhibited significantly higher peptide yields in aged than in young posterior and lateral IVD and also in female plaque compared to control artery. This segment coincides with an HC1 fragment which can be released during beta thrombin activation. Similarly, one segment corresponding to the fibronectin type-III (Fn) 8 domain of COL14A1 **(C)** also yielded the same significantly higher peptides in aged anterior IVD than in young as seen for male plaque when compared to control artery. Apolipoprotein B **(D)** contained a single segment near the centre of the protein which exhibited significantly higher peptide yields in young anterior IVD than aged and also in male control artery compared to plaque. This segment lies within the fourth heparin-binding domain (HB) of the protein.

The higher peptide yields observed within the MG2 domain of A2M for all three young IVD OAF tissue regions compared to aged was discussed earlier **(Fig. 4B)**. Interestingly, the same equivalent pattern was observed in control artery when compared to plaque for both male and female groups **(Fig. 13A)**. Since this domain frames the entrance to the cavity of A2M [64], which functions to entrap proteinases within its dimeric cage [63], differences within it may reflect its conformational state (bound or unbound). As such, it is possible that an equivalent change in state may exist in aged IVD as in arterial plaque as a consequence of ageing and age-related atherosclerosis, which may indicate a dynamic shift in protease inhibition within both tissues.

Thrombin is a serine protease whose main function is to cleave fibrinogen into fibrin during the process of coagulation [121]. This is achieved via activation by prothrombinases (e.g. factor Xa), which cleave prothrombin both at the N-terminal end of its light chain (LC; yielding fragments [F] 1 and 2) and between its light and heavy chain. These chains are then held together by a single disulphide bridge, to form the active α-thrombin [121, 122]. One segment near the N-terminal end of prothrombin’s heavy chain yielded significantly more peptides for aged posterior and lateral IVD than young **(Fig. 13B)**. This pattern was also observed for female arterial plaque when compared to control, potentially indicating a mechanism or consequence that may be conserved between IVD ageing and arterial atherosclerosis. α-thrombin can undergo degradation into two less active forms, β- and γ-thrombin, both of which yield an HC1 fragment [121, 122]. Interestingly, the conserved, significant differences in peptide yields observed for both IVD ageing and arterial atherosclerosis fall specifically within the protein region responsible for yielding this fragment. Should the rise in peptide yields indicate a higher presence of this fragment, it could be indicative of thrombin degradation in both aged and atherosclerotic tissues. Age-associate differences were observed in both fibrinogen **(Fig. 4C)** and A2M **(Fig. 13A)**. Since thrombin functions to activate fibrinogen and A2M is capable of regulating fibrin degradation [67], structure-associated alterations in these three proteins may reflect a change in the progress or state of coagulation, which is present in both ageing and atherosclerosis.

Collagen XIV belongs to the subfamily of so-called fibril-associated collagens with interrupted triple-helices (FACITs). It exists as a three-pronged homotrimer of COL14A1 chains which function to bridge adjacent collagen I fibrils [123]. Collagen XIV was found to prevent the lateral fusion of collagen I fibrils by limiting fibril diameter [124] and is therefore highly present in tissue regions bearing high mechanical stress [125]. Previous analysis of the IVD ageing dataset showed a lower relative abundance of COL14A1 in full aged IVD compared to young [31] which may contribute to the remodelling of fibrillar collagen. Here, we show that COL14A1 from aged anterior OAF yields more peptides than young within the Fn8 domain, near the centre of the of its structure **(Fig. 13C)**. Interestingly this same domain also yielded higher peptides in male atherosclerotic plaque than in control artery, perhaps indicating a similar structure-associated consequence as observed in aged IVD. The AF is comprised of hierarchical, concentric rings of collagen fibrils (lamellae), which are thought to be remodelled in ageing [126]. Similarly, atherosclerotic plaques also contain collagen fibrils which are synthesised by smooth muscle cells during disease progression, and their degradation is thought to affect plaque stability during disease [112]. It is possible that structure-dependent changes in collagen XIV may contribute to the degradation or remodelling of collagen in both tissues, through the disruption of the interfibrillar bridges they form.

APOB is the primary lipoprotein component of LDLs (and very low-density lipoproteins, VLDLs) and through its tight association, makes up ∼50% of the of the surface of these spherical particles. APOB is highly implicated in atherosclerosis, through its interaction with cell surface LDL-receptors which drive the accumulation of oxLDLs in the intima, under the vascular epithelium [119]. Although the association of APOB in IVD ageing remains elusive, a recent study showed that levels of oxLDLs increase during IVD degeneration (particularly in the nucleus pulposus and OAF) [120]. Another demonstrated that patients with IVD herniation exhibited raised serum levels of APOB compared to healthy controls [127]. Six heparin-binding (HB) domains exist along the structure of this lipoprotein which are thought to interact with proteoglycans within the subendothelium during early atherosclerosis [128]. Interestingly, similar raised peptide yields were observed in segments within the HB4 domain of APOB from both control male artery compared to plaque and from young IVD OAF compared to aged **(Fig. 13D)**. Disruptions or changes associated to this domain could have implications on the ability of LDLs to interact with proteoglycans within these tissues, potentially providing evidence of shared consequences to both ageing and atherosclerosis.

A2M, prothrombin, collagen XIV and APOB exhibited regional peptide yield differences with possible shared structural consequences between ageing in IVD and atherosclerosis in artery. As such, these newly identified, common markers may provide new links between ageing and atherosclerosis, albeit in two disparate tissues. These potentially corroborate the notion that age-related diseases harbour accelerated forms of tissue ageing, with mechanisms and consequences that may be comparable in both.

## Conclusion

The identification of age-susceptible protein structures that are conserved between species, organs and in age-related disease, and the characterisation of shared mechanisms and functional consequences, is crucial for the understanding of connective tissue ageing. The application of PLF as a proteomic discovery tool to ageing human IVD, ageing mouse lung and human arterial atherosclerosis datasets provided potentially crucial evidence of common age-associated differences to protein structures, some of which were conserved between species, organs and in age-related disease. Crucially, PLF identifies structure-associated differences in ECM proteins which remain undetected by conventional whole protein quantification approaches **(Fig. 14A)**, therefore enabling a more complete assessment of tissue proteostasis in ageing and disease than previously achieved. Furthermore, the potential of PLF to be used for the interrogation of functional consequences and mechanisms of ageing and disease is particularly promising; with possible changes in macromolecular structure, enzyme/inhibitor activity, protein activation and ECM-cell communication being revealed **(Fig. 14B)**. Finally, the prospect that ECM components and their associated proteins may be subjected to potentially similar mechanisms or consequences of ageing in both mouse and human and in age-related atherosclerosis is interesting, as it suggests the potential for universal targets that may be present irrespective of differences in lifespans, tissue functions and disease progressions.

**Figure 14.**
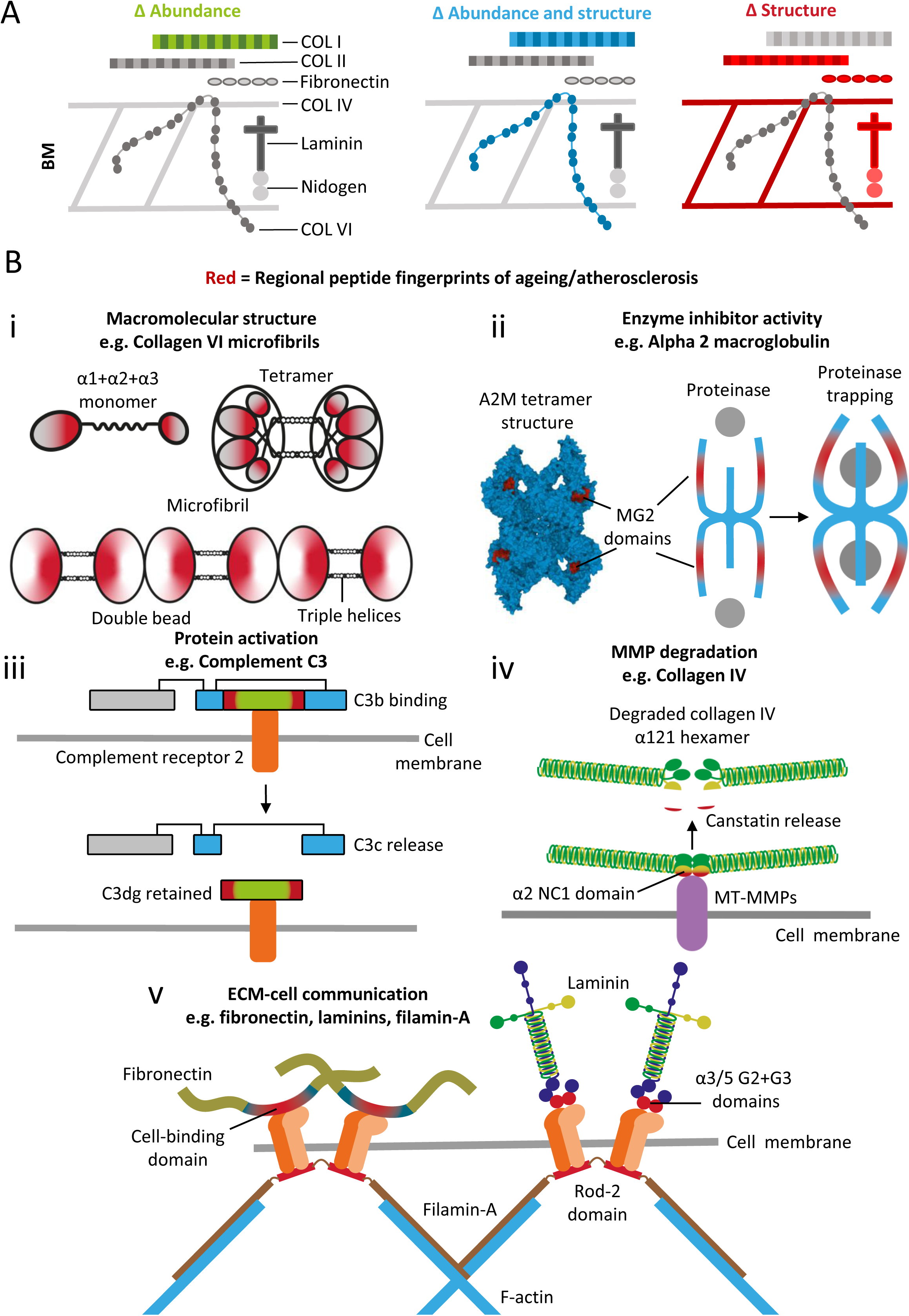
PLF identifies targets of ageing and atherosclerosis that are unique to the methodology, alongside potentially affected mechanisms and functional consequences. For instance, PLF revealed several basement membrane components in the IVD that were sensitive to structure-associated differences, but did not change in relative abundance (as reported previously [31]) between aged and young **(A)**. Multiple underlying mechanisms or functional consequences of ageing/atherosclerosis were potentially revealed through regional peptide fingerprints (significant differences) within protein structures **(B)**. PLF identified possible changes in: **(i)** macromolecular structure, such as for the collagen VI microfibril, where globular regions within its double bead, closest to the triple helices, exhibited differences in peptide yield in IVD ageing (for α1 and α3) and arterial atherosclerosis (for α2); **(ii)** enzyme inhibition activity, such as for A2M where the MG2 domains that frame the entrance to the proteinase entrapping cavity exhibited peptide yield differences which may be linked to distinct conformational states between aged and young IVD and atherosclerotic and control artery; **(iii)** protein activation, such as for complement C3 where regions corresponding to both ends of the C3dg fragment exhibited significant differences in peptide yield, indicating the heightened potential transition of C3b to C3c in young compared to aged IVD; **(iv)** MMP degradation, such as for collagen IV where the NC1 domain of the α2 chain yielded significantly more peptides in young mouse lung than in aged, which may be linked to the known degradation and release of the canstatin matrikine by MT-MMPs; **(v)** ECM-cell communication, such as a perturbed integrin-mediated link between the rod-2 domain of the cytoskeletal filamin-A and the cell-binding domain of fibronectin or G2 and G3 domains of laminin, all of which exhibited significant differences in peptide yield in both human IVD and mouse lung ageing.

## Materials and Methods

### Dataset sourcing and summaries of sample preparation and mass spectrometry methods

All label-free proteomic LC-MS/MS datasets used in this study were sourced from the Proteomics Identification Database (PRIDE) repository.

The young and aged human IVD datasets were generated by Tam and Chen *et al.* and originally used in the development of the spatiotemporal IVD proteomic resource - DIPPER (http://www.sbms.hku.hk/dclab/DIPPER/), already published [31]. As detailed in their original publication, 11 separate IVD regions were carefully dissected from three discs (L3/4, L4/5 and L5/S1), acquired from one aged (59 yr old) and one young (16 yr old) male. Only three of these regions were analysed for this study (posterior, left lateral and anterior portions of the OAF). To summarise their procedures pertinent to this study, after pulverisation from frozen with a freezer mill, proteins were mechanically and chemically extracted by ten freeze-thaw cycles and sonication prior to agitation for two days within an extraction buffer comprised of guanidine hydrochloride, sodium acetate, 6-aminocaproic acid and a protease inhibitor cocktail. Samples were then ultracentrifuged and supernatants were ethanol-precipitated prior to re-centrifugation to leave protein pellets. These were re-solubilised by sonication in urea and ammonium bicarbonate buffer prior to reduction with TCEP and alkylation with iodoacetamide. Samples were then digested with trypsin/|LysC and peptides then splt into four fractions by high pH reversed phase fractionation. LC-MS/MS datasets were generated by data-dependant acquisition using an Orbitrap Fusion Lumos Tribrid Mass Spectrometer. Raw MS datasets were downloaded from PRIDE - Project ID PXD017740 for PLF analysis. For more detailed methods of tissue sourcing, sample preparation and MS, please refer to the original publication [31].

The young and aged mouse lung datasets were originally generated by Angelidis and Simon *et. al.* and used in the investigation of ageing by deep tissue quantitative proteomics, already published [30]. To summarise their procedures pertinent to this study, whole bulk lung was taken from young and aged mice (3 and 24 months old; N = 4) and homogenised in PBS containing protease inhibitors. After centrifugation, soluble proteins were separated from insoluble via three incubation steps with three separate buffers of increasing chaotropic activity. All three buffers contained NaCl, Tris-HCl, glycerol, protease inhibitors, Benzonase (Merck) and IGEPAL-CA- 630 (Sigma). However, buffer 2 contained added 0.5% sodium deoxycholate and 0.1% SDS while buffer 3 had higher concentrations of salt (500 mM NaCl compared to 150 mM), sodium deoxycholate (2%) and SDS (1%).

Treatment with these buffers resulted in an insoluble protein pellet, enriched in ECM proteins, which was further heated, reduced in TCEP and alkylated in chloroacetamide within a guanidinium hydrochloride buffer. This was then mechanically disrupted using a micro-dounce prior to ultrasonication and overnight digestion with trypsin/LysC to create an ECM-rich peptide fraction. Peptides were purified and LC-MS/MS datasets generated by data-dependant acquisition using a Q-Exactive Hybrid Quadrupole-Orbitrap Mass Spectrometer (Thermo). Raw MS datasets was downloaded from PRIDE - Project ID PXD012307 for PLF analysis. For more detailed methods of tissue sourcing, sample preparation and MS, please refer to the original publication [30].

The human atherosclerotic artery datasets were originally generated by Liang and Ward *et al.* and used in the proteomic investigation of carotid plaques in men and women, already published [32, 33]. To summarise their procedures pertinent to this study, biopsies (4 mm) were taken from atherosclerotic plaque centre regions and internal control regions of carotid arteries obtained from patients (N = 10 male and 10 female; age range 60 to 83 yrs) undergoing endarterectomy. These were flash frozen and crushed prior to sonication in TriZol LS (Life Technologies). Phase separation with chloroform was performed to remove nucleic acids and proteins precipitated by isopropyl alcohol. The centrifuged protein pellet was then resuspended in a urea/thiourea buffer containing protease inhibitors (PefaBloc; Sigma) prior to reduction with dithiothreitol and alkylation with iodoacetamide. Samples were filtered (3 kDa cut-off) and digested with trypsin. Peptides were purified and LC-MS/MS datasets generated using an LTQ Orbitrap Velos Pro Mass Spectrometer (Thermo). Raw MS datasets was downloaded from PRIDE - Project ID PXD003930 for PLF analysis. For more detailed methods of tissue sourcing, sample preparation and MS, please refer to the original publication [32].

### Peptide identification

RawConverter (Scripps Research Institute; CA, USA) [129] was used to convert all raw MS spectrum files to Mascot MGF files prior to peak spectra searches against the Uniprot database (Swiss-Prot and TreEMBL; 2018) [130]. MS/MS ion searches were performed using Mascot (Matrix Science, MA, USA) with the following parameters. Human IVD datasets – taxonomy: *homo sapiens* (161,629 proteins and isoforms), fragment and parent tolerances: 20 ppm and 6 ppm respectively (monoisotopic), fixed modifications: +57 Da on cysteine (carbamidomethylation), variable modification: +16 Da on methionine (oxidation), enzyme: trypsin, max missed cleavages: 2, peptide charge: 2+, 3+. Mouse lung datasets – taxonomy: *mus musculus* (84,416 proteins and isoforms), fragment and parent tolerances: 20 ppm and 4.5 ppm respectively (monoisotopic), fixed modification: +57 Da on cysteine (carbamidomethylation), variable modification: +16 Da on methionine and proline (oxidation), enzyme: trypsin, max missed cleavages: 2, peptide charge 2+, 3+. Human artery datasets – taxonomy: *homo sapiens* (161,629 proteins and isoforms), fragment and parent tolerances: 0.6 Da and 10 ppm respectively (monoisotopic), fixed modifications: +57 Da on cysteine (carbamidomethylation), variable modification: +16 Da on methionine (oxidation), enzyme: trypsin, max missed cleavages: 2, peptide charge: 2+, 3+.

Exported Mascot search results (DAT files) were then imported into Scaffold 5 (Proteome Software; OR, USA). For the human IVD datasets, pH fractionated peptide sample datasets were combined using MudPIT (Stowers Institute; MO, USA) [131] to produce a single dataset per IVD region (posterior, lateral and anterior). High confidence peptide spectrum matches were identified by standard LFDR scoring. Peptide false discovery rates (FDR) were calculated within Scaffold 5 using the PeptideProphet algorithm (WA, USA) [132] based on peptide probabilities assigned via the Trans Proteomic Pipeline. Peptides exclusive to their matched proteins were thresholded to 95% peptide probability minimum for all datasets, resulting in low FDRs of 0.7% for the human IVD dataset, 1.5% for the mouse lung dataset and 1.4% for the human artery dataset. Peptide lists were exported from Scaffold 5 as CSV files which were used for PLF analysis.

### Peptide Location Fingerprinting

Peptide location fingerprinting was applied using our predeveloped MPLF webtool as previously described (https://www.manchesterproteome.manchester.ac.uk/#/MPLF) [26, 27]. Analyses were performed separately on each OAF region (posterior, lateral and anterior) for the human IVD young vs. aged datasets and also separately on males and females for the human arterial atherosclerotic plaque vs. control datasets.

Peptide list CSV files were imported into the webtool, and the primary sequences of matched proteins were bioinformatically divided into 50 aa-sized segments. Identified peptide sequences were then mapped and quantified within each segment by spectral count. Peptide sequences which spanned two connecting segments were counted in both. Total spectral counts were then summed per segment. To detect protein region-specific differences in peptide yield that were unskewed by differences in whole protein abundance, summed peptide counts per segment were median normalised based on the experiment-wide total spectrum counts of their corresponding whole proteins. Additionally, proteins which were exclusively present in one experimental group (young or control) but not the other (aged or plaque) were excluded from comparisons. These steps minimised the effect of whole protein relative abundance on regional comparisons of peptide yield across protein structure and ensured that the identification of proteins with structure-related differences were independent of protein presence or quantity. Modular fluctuations in peptide yield across protein structures were revealed by subtracting the average, normalised peptide counts per segment in one experimental group from the other and dividing them by the segment length (50 aa). Average, normalised peptide counts in each segment were statistically compared between experimental groups using Bonferroni-corrected, repeated measures ANOVAs (unpaired for young vs aged; paired for plaque vs control). For further details on PLF and its application, please refer to our MPLF webtool development publication [26].

## Author Contributions

AE conceived and designed the study, performed all PLF analysis, interpreted the data, prepared the figures, and wrote the paper. AE and MJS contributed to the acquisition of funding. MO contributed to the running of PLF and to the maintenance, support, and continued development of the MPLF webtool. MJS and AT contributed to study conception. MJS contributed to experimental design and preparation of figures. PC, VT, LJW, JAH, XMY, HBS, DC and MJS contributed to the interpretation of results. All authors contributed to editing of the paper.

## Funding Disclosure

This study was jointly funded by the Manchester Institute for Collaborative Research on Ageing (MICRA) (AE [PI], AT, MO and MJS [Co-I]) and Walgreens Boots Alliance (WBA) (MJS), UK. In line with WBA’s Corporate Social Responsibility Policies, no animal testing has been conducted as part of this study. All mouse mass spectrometry datasets have been acquired (downloaded) from a public repository - PRIDE. MO is supported by the Wellcome Trust (206194), DC is supported by The Research Council of Hong Kong (E-HKU703/18) and PC by the Shenzhen Health Commission (SZHC) “Key Medical Discipline Construction Fund” (SZXK077). JAH and MJS would like to acknowledge funding from the UK Engineering and Physical Sciences Research Council (EPSRC; EP/V011065/1). The work on the human atherosclerotic artery datasets was supported by grants to XMY from the Swedish Heart Lung Foundation, Stroke Foundation, Olle Engkvist Foundation, Swedish Gamla Tjänarinnor Foundation and Linköping University Hospital Research Fund.

## Data Availability

The LC-MS/MS datasets analysed by PLF in this paper were downloaded from the PRIDE repository: human IVD ageing - PXD017740; mouse lung ageing - PXD012307; human arterial atherosclerosis - PXD003930. PLF analyses of all datasets, and associated protein schematics will be interactively viewable and downloadable via the MPLF webtool at https://www.manchesterproteome.manchester.ac.uk/#/MPLF under the “Location Fingerprinter” tab [26, 27] upon journal acceptance.

## Conflict of Interest

The authors declare no conflicts of interest. WBA approved manuscript submission but exerted no editorial control.

## Supporting information

Table S1_Peptide Report for Anterior OAF

Table S2_Peptide Report for Left Lateral OAF

Table S3_Peptide Report for Posterior OAF

Table S4_PLF of Young vs Aged Human Anterior OAF

Table S5_PLF of Young vs Aged Human Left Lateral OAF

Table S6_PLF of Young vs Aged Human Posterior OAF

Table S7_Peptide Report for Mouse Lung

Table S8_PLF of Young vs Aged Mouse Lung

Table S9_Peptide Report for Male Human Artery

Table S10_Peptide Report for Female Human Artery

Table S11_PLF of Control vs Plaque Male Human Artery

Table S12_PLF of Control vs Plaque Female Human Artery

## Notes

https://www.manchesterproteome.manchester.ac.uk/#/MPLF

http://proteomecentral.proteomexchange.org/cgi/GetDataset?ID=PXD017740

http://proteomecentral.proteomexchange.org/cgi/GetDataset?ID=PXD012307

http://proteomecentral.proteomexchange.org/cgi/GetDataset?ID=PXD003930

### Competing Interest Statement

The authors have declared no competing interest.

