## Supporting Figures for "Peptide location fingerprinting identifies species- and tissue-conserved structural remodelling of proteins as a consequence of ageing and disease"

Protein presence – IVD OAF ageing

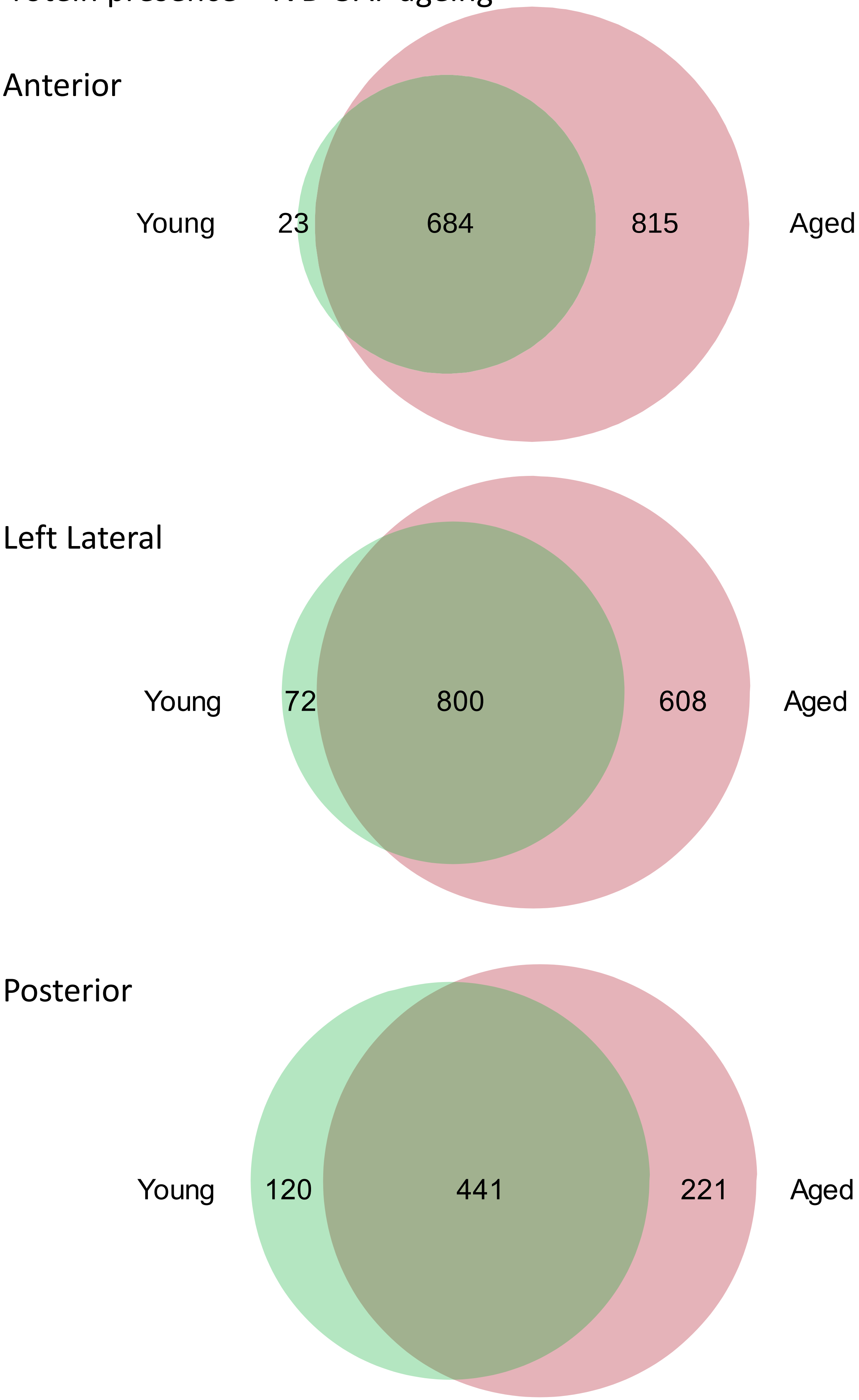

### PCA of peptide spectrum matches – IVD OAF ageing

#### Anterior

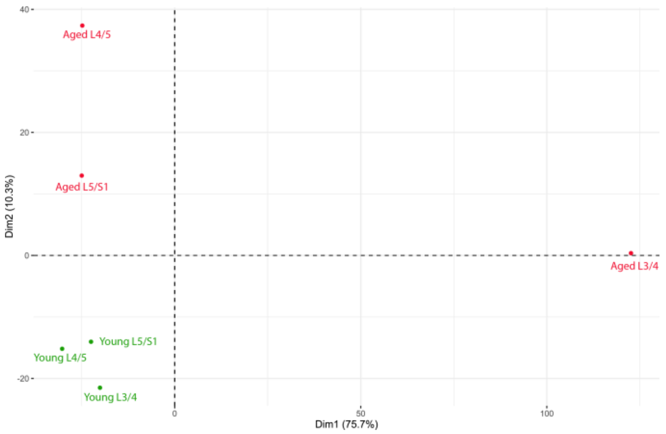


#### Left Lateral

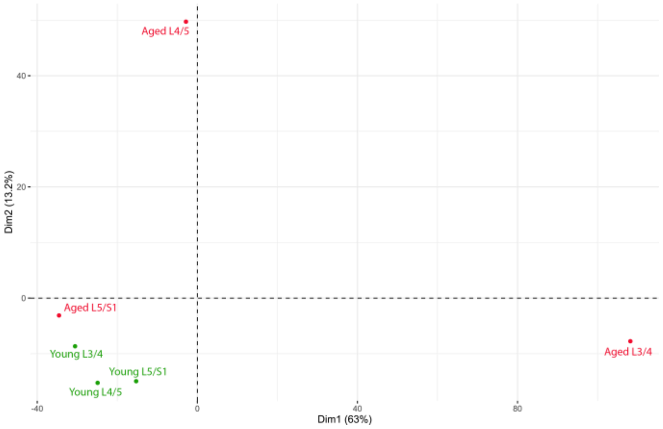


#### Posterior

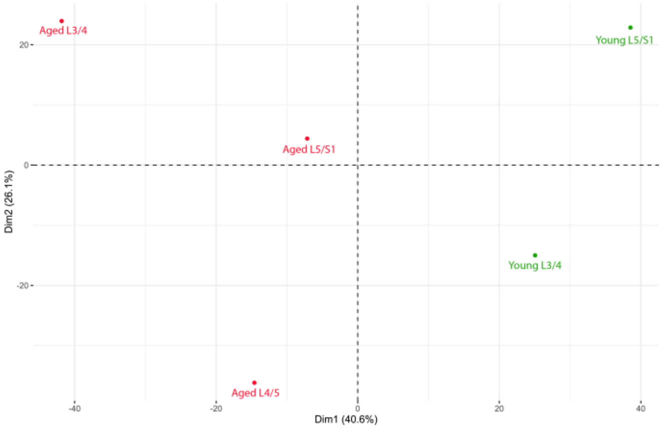

### Biomarker candidates shortlisted – IVD OAF ageing

**Shared Biomarker Candidates**

A2M, ABI3BP, AHNAK, C3, C4A, CILP, CILP2, COL11A2, COL3A1, COL6A1, COL6A3, COL8A1, COMP, CSPG4, FBLN1, FLNA, FNDC1, HSPA1A, HSPA1B, HTRA1, LAMB2, LRP1, MATN2, PLEC, TIMP1, TIMP3, VCAN

**Anterior-only Biomarker Candidates**

A1BG, ACTB, ACTN1, ACTN4, ANGPTL5, ANXA7, APCS, APOA4, APOB, ARF1, ARF3, ATP1A1, C1S, C6, CA2, CALD1, CAST, CAVIN3, CD109, CDH13, CHAD, CHRDL2, COL14A1, COL4A2, COL5A2, CPNE3, CPXM2, DKK3, DPYSL2, ECM1, EEF1G, EMILIN1, EMILIN3, ENO3, ENPP2, ERP44, FLNB, FTL, H1-0, H2BC11, H2BC21, H2BC3, HBD, HEXA, HHIPL2, HNRNPK, HP, IGL@, ITGB1, ITIH1, KRT1, KRT10, KRT19, LAMB1, LRG1, LTA4H, MMP2, MRC2, MXRA5, MYH10, MYH11, NACA, NID2, OLFML1, ORM1, PAM, PDGFC, PDIA6, PEBP1, PFN1, PLG, PRKCSH, PSMA7, PYGL, PYGM, RBMX, RPN1, SERPINA1, SERPINA3, SERPINB6, SERPIND1, SERPINF1, SERPING1, SET, SNC73, SOD3, SPTBN1, SRPX2, TGM2, THBS2, THBS4, TINAGL1, VASN, XYLT1

**Anterior and Left Lateral Shared Biomarker Candidates**

CLEC3B, COL15A1, COL2A1, COL9A2, ECM2, FRZB, HAPLN3, HNRNPU, ISLR, ITIH2, LAMA2, LAMA5, MELTF, OAF, P4HA1

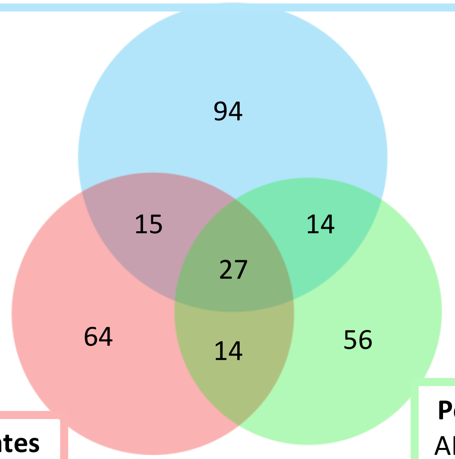


**Anterior and Posterior Shared Biomarker Candidates**

C2, CFH, FBN1, FGA, FGB, FN1, H4C1, HPX, IGLC1, NUCB2, TNC, TTR, TUBA1B, VTN

**Left Lateral-only Biomarker Candidates**

ACLY, ACTN3, ARHGDIB, CALU, CCN2, CDH1, COL5A1, CTGF, CTSG, DPT, DYNC1H1, EIF4A2, ENPP1, FLNC, GAA, GANAB, GAPDH, GBE1, GCS1, GSTP1, H1-5, HIST1H1E, HNRNPA3, HNRNPH1, HSP90AB1, HTRA3, IDH2, IGFBP7, IGHM, ITIH4, ITIH6, KRT9, LAMC1, LCP1, LGALS3BP, LOXL3, MAP1B, MDH2, MOGS, MYOM2, NID1, NNMT, PGK1, PLS3, PRDX3, PRKAR1A, PRTN3, PXYLP1, RAP1B, RPN2, S100A9, SCIN, SEMA3C, SEMA3E, SERPINA4, SLC4A1, SPARC, SPTA1, SRI, THBS1, TKT, TUBB4B, UGP2, VWA1

**Left Lateral and Posterior Shared Biomarker Candidates**

ACAN, CCDC80, CLTC, COL11A1, F2, FHL1, LTBP2, PPIC, PRG4, RACK1, SMOC2, TNXB, UGDH, VIM

**Posterior-only Biomarker Candidates**

AKR1A1, AMBP, ANGPTL2, ANXA2, ANXA5, APOA1, APOE, BGN, C9, CALM1, CALM2, CALM3, CAPS, CKM, CLEC3A, COL1A1, COL6A2, COL9A1, CRYAB, DF, EEF1A1, EEF1A1P5, EZR, F13A1, FBXO2, FSCN1, GC, GDI2, GLUD1, GSN, TF, HRG, HSPA5, HSPA8, HSPG2, IGK@, KNG1, LOX, MSN, NCL, NUCB1, ORM2, PCOLCE, PCOLCE2, PDCD6IP, PGD, PLOD1, PLTP, PTRF, PXDN, RHOA, SERPINE1, SMOC1, THBS3, TUBA1A, TXNDC5

#### Protein presence in ECM Fraction – Mouse Lung ageing

A

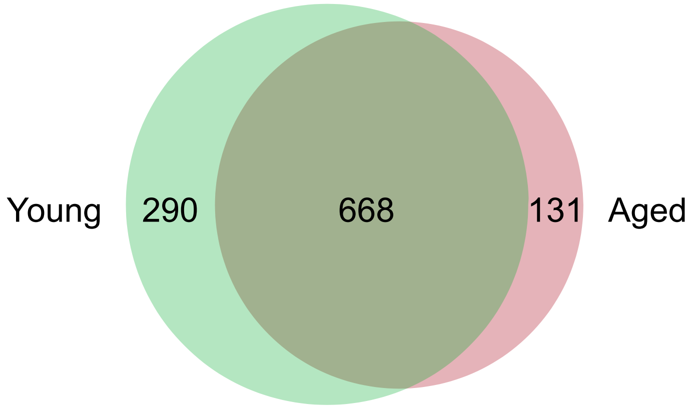


#### PCA of peptide spectrum matches – Mouse Lung ageing

B

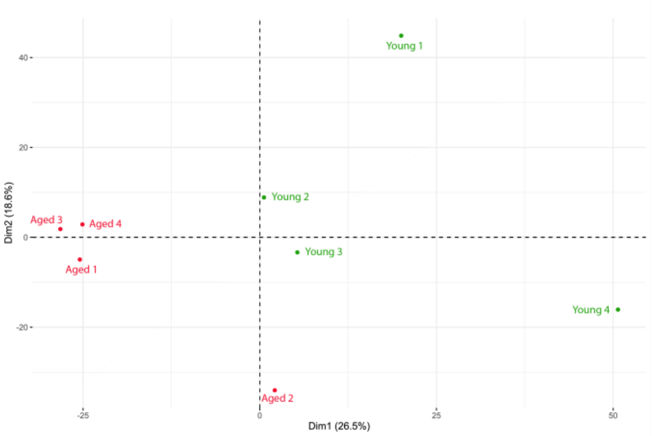

### Shared and unique ageing biomarker candidates between mouse Lung and human

**Shared Biomarker Candidates between Human IVD and Mouse Lung Ageing**

ABI3BP, AHNAK, APOA1, ATP1A1, CFH, COL15A1,COL1A1, COL4A2, COL6A2, DYNC1H1,ECM1, EZR, FBN1, FGA, FGB, FLNA, FN1, GDI2, HSPA5, HSPG2, IGFBP7, KRT1, LAMA2, LAMA5, LAMB1, LAMB2, LAMC1, MYH10, MYOM2, NCL, PDCD6IP,PXDN, SEMA3E, TGM2, TINAGL1,TXNDC5, VTN

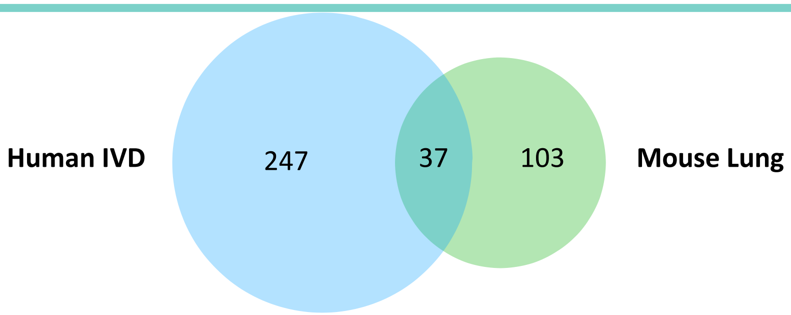


**Ageing Human IVD-only Biomarker Candidates**

A1BG, A2M, ACAN, ACLY, ACTB, ACTN1, ACTN3, ACTN4, AKR1A1, AMBP, ANGPTL2, ANGPTL5, ANXA2, ANXA5, ANXA7, APCS, APOA4, APOB, APOE, ARF1, ARF3, ARHGDIB, BGN, C1S, C2, C3, C4A, C6, C9, CA2, CALD1, CALM1, CALM2, CALM3, CALU, CAPS, CAST, CAVIN3, CCDC80, CCN2, CD109, CDH1, CDH13, CHAD, CHRDL2, CILP, CILP2, CKM, CLEC3A, CLEC3B, CLTC, COL11A1, COL11A2, COL14A1, COL2A1, COL3A1, COL5A1, COL5A2, COL6A1, COL6A3, COL8A1, COL9A1, COL9A2, COMP, CPNE3, CPXM2, CRYAB, CSPG4, CTGF, CTSG, DF, DKK3, DPT, DPYSL2, ECM2, EEF1A1, EEF1A1P5, EEF1G, EIF4A2, EMILIN1, EMILIN3, ENO3, ENPP1, ENPP2, ERP44, F13A1, F2, FBLN1, FBXO2, FHL1, FLNB, FLNC, FNDC1, FRZB, FSCN1, FTL, GAA, GANAB, GAPDH, GBE1, GC, GCS1, GLUD1, GSN, GSTP1, H1-0, H1-5, H2BC11, H2BC21, H2BC3, H4C1, HAPLN3, HBD, HEXA, HHIPL2, HIST1H1E, HNRNPA3, HNRNPH1, HNRNPK, HNRNPU, HP, HPX, HRG, HSP90AB1, HSPA1A, HSPA1B, HSPA8, HTRA1, HTRA3, IDH2, IGHM, IGK@, IGL@, IGLC1, ISLR, ITGB1, ITIH1, ITIH2, ITIH4, ITIH6, KNG1, KRT10, KRT19, KRT9, LCP1, LGALS3BP, LOX, LOXL3, LRG1, LRP1, LTA4H, LTBP2, MAP1B, MATN2, MDH2, MELTF, MMP2, MOGS, MRC2, MSN, MXRA5, MYH11, NACA, NID1, NID2, NNMT, NUCB1, NUCB2, OAF, OLFML1, ORM1, ORM2, P4HA1, PAM, PCOLCE, PCOLCE2, PDGFC, PDIA6, PEBP1, PFN1, PGD, PGK1, PLEC, PLG, PLOD1, PLS3, PLTP, PPIC, PRDX3, PRG4, PRKAR1A, PRKCSH, PRTN3, PSMA7, PTRF, PXYLP1, PYGL, PYGM, RACK1, RAP1B, RBMX, RHOA, RPN1, RPN2, S100A9, SCIN, SEMA3C, SERPINA1, SERPINA3, SERPINA4, SERPINB6, SERPIND1, SERPINE1, SERPINF1, SERPING1, SET, SLC4A1, SMOC1, SMOC2, SNC73, SOD3, SPARC, SPTA1, SPTBN1, SRI, SRPX2, TF, THBS1, THBS2, THBS3, THBS4, TIMP1, TIMP3, TKT, TNC, TNXB, TTR, TUBA1A, TUBA1B, TUBB4B, UGDH, UGP2, VASN, VCAN, VIM, VWA1, XYLT1

**Ageing Mouse Lung-only Biomarker Candidates**

Cavin1, Cep135, Col4a3, Col4a5, Crocc2, Dsp, Eln, Fermt2, Lama3, Lama4, Lamc2, Macroh2a1, Mprip, Myl6, Myo1b, Myo1c, Postn, Serpina1d, Serpina3k, Serpinc1, Serpinh1, Tgfbi, Ttn, Agrn, Alb, Atp5f1b, C4b, Ddx5, Efemp1, Ehd4, Hspa12b, Pakap, Poc5, Rpl7a, Serpina1b, Tubb5, Ccdc170, Ccdc187, Fbln5, Itga8, Mylk, Ppp2r1a, Slc25a4, Tjp1, Top2b, Vwf, Aco2, Actr2, Add1, Ap2a2, Arhgef7, Arpc1b, Ces1d, Cma1, Coro1c, Cp, Dag1, Dhx9, Dpep1, Dstn, Emid1, Eppk1, Flii, H1-2, Hadhb, Hmcn1, Hnrnpl, Hpgd, Ighg2b, Khdrbs1, Kpnb1, Lama1, Lsp1, Macf1, Mlf1, Mmrn1, Myh2, Myh7, Myh8, Myo18a, Parp1, Pdia3, Pi15, Plcb3, Ppia, Ppp1cc, Prx, Rac1, Rpl14, Ruvbl1, Samhd1, Sec31a, Smc3, Sorbs1, Spata6, Sptan1, Sqor, Stab1, Svep1, Syne1, Tns3, Tpm1, Wdr1

Protein presence – Artery atherosclerosis

Male

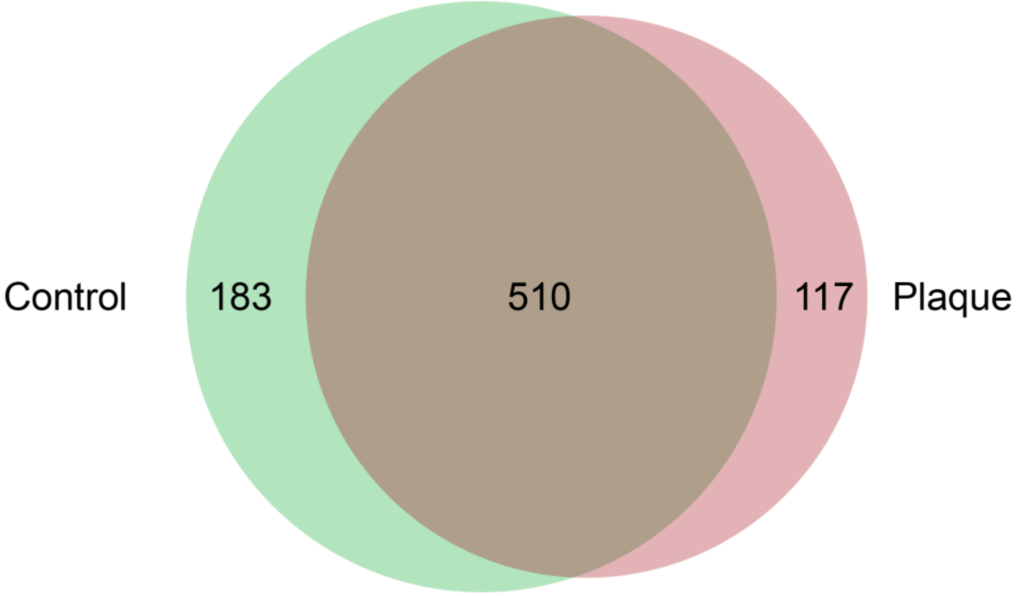


Female

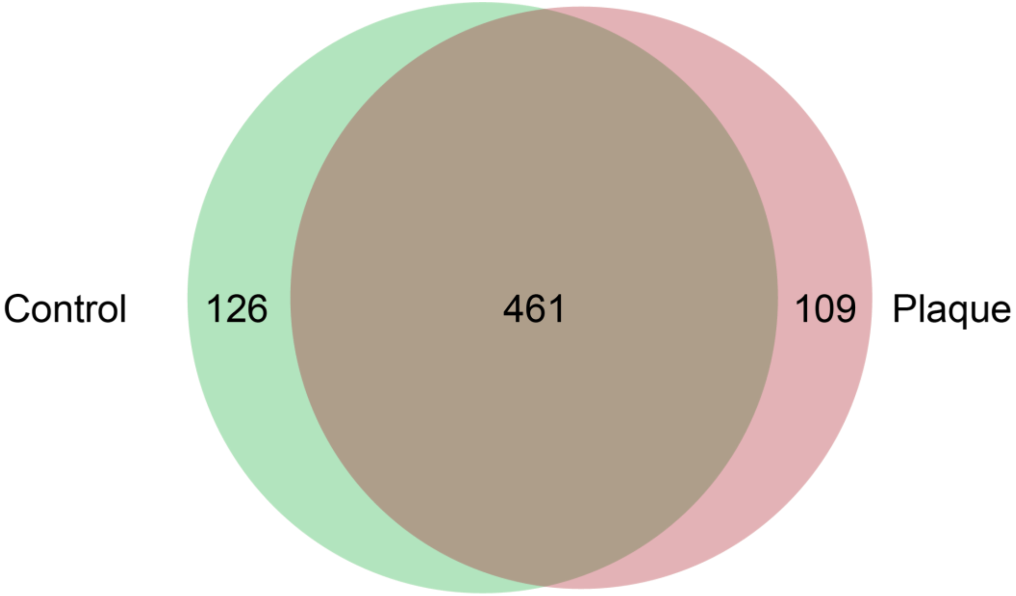

### PCA of peptide spectrum matches – Artery atherosclerosis

#### Male

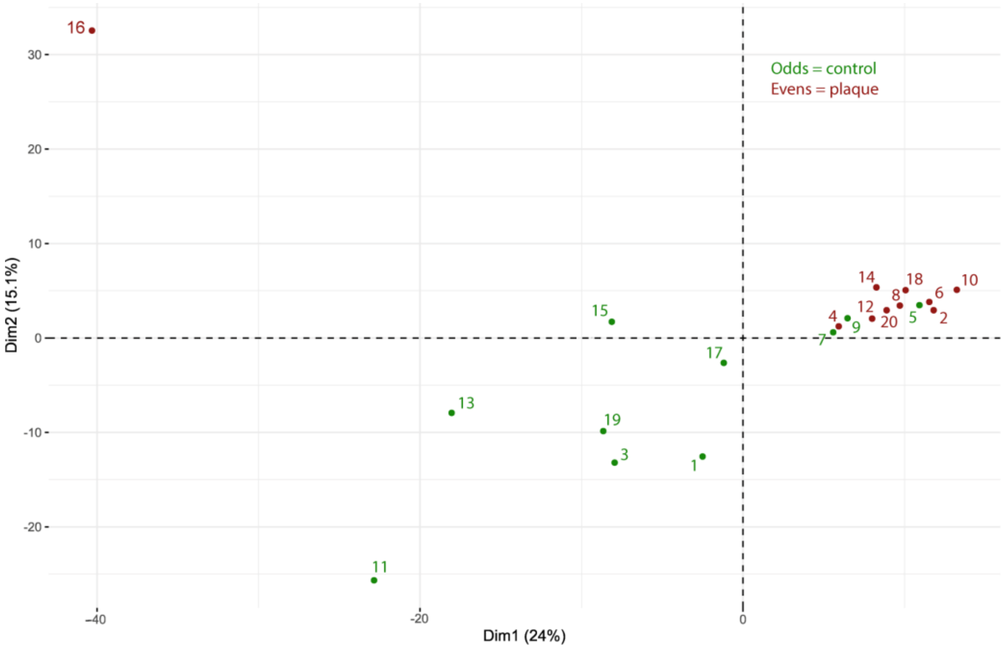


#### Female

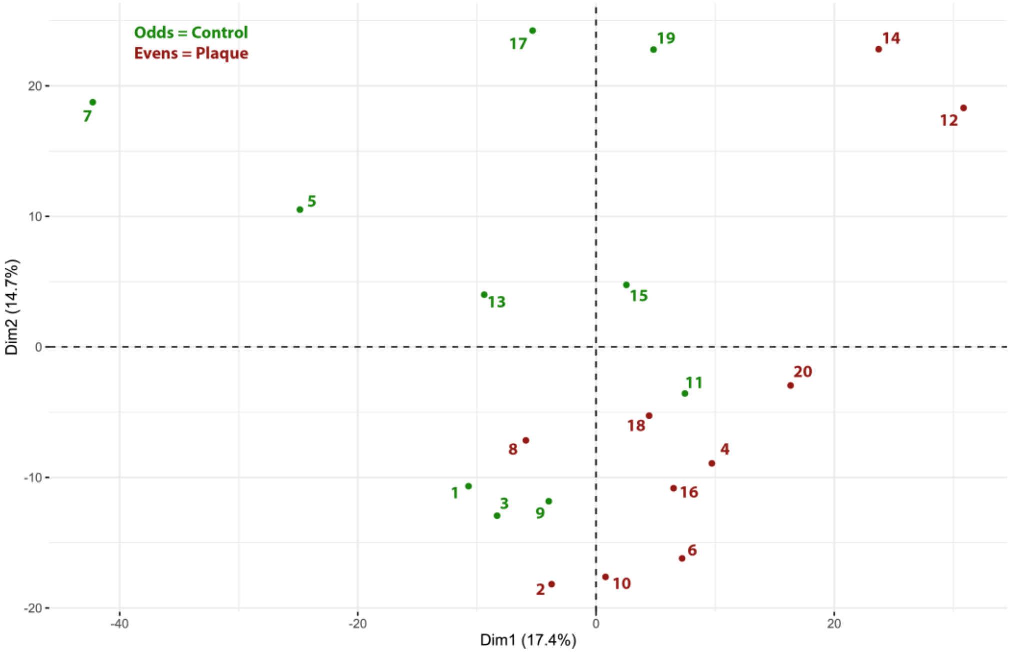

### Biomarker candidates shortlisted - Artery atherosclerosis

**Shared Biomarker Candidates**

ACTC1, ACTN4, AHNAK, ANK1, ANXA1, AOC3, APOB, APOL1, APOM, AQP1, C3, C4B, C7, C8B, CAVIN1, CDH13, CFI, COL12A1, COL18A1, COL4A2, COL6A2, CSRP1, CTSB, DPYSL3, EEF2, FBLN1, FCN3, FERMT3, FLNA, FMOD, FTH1, GM10881, GPNMB, GSTO1, HSP90AB1, HSPG2, IQGAP1, LAMC1, LAMP1, LASP1, LBP, LRP1, LTBP2, MMP12, MYH11, MYH9, OGN, PFN1, PGD, PLEC, RBP4, RTN4, SERPINA4, SERPIND1, SOD3, SPTA1, SPTB, TF, THBS1, TIMP1, TLN1, TNC, TNXB, YWHAG

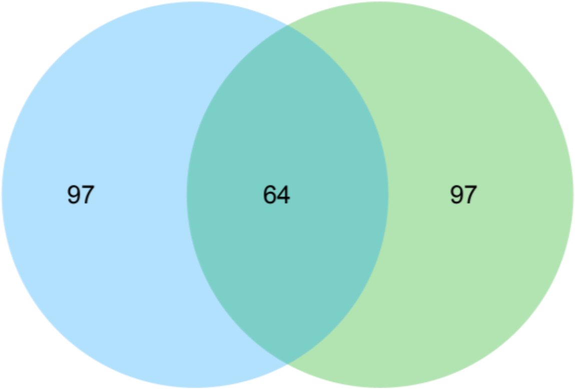


**Female-only Biomarker Candidates**

A2M, ACTR2, ACTR3, AEBP1, AFM, AGT, AMBP, APOA1, APOE, ARHGDIB, ARPC2, ASPN, C1QA, C1QB, C2, C6, C9, CALU, CD14, CD36, CD44, CFHR1, CFL1, CLIC1, CLTC, CNDP2, CNN1, COL1A2, COL4A1, COL6A1, COL6A3, CORO1A, CPN2, CRIP2, CYB5R3, DEFA1, DEFA3, EFEMP1, EIF4A1, EMILIN2, F10, F12, F13A1, F2, FBLN5, FGG, FLJ00385, FN1, GAPDH, GAS6, H3-3A, H4C1, HPR, HSPA1A, HSPA1B, HTRA1, IGFALS, IGHG3, ITGB1, ITGB2, ITIH4, KNG1, LTBP1, MYH10, MYL12A, MYL9, NID1, PDIA3, PLD3, PLS3, PLTP, PON1, POSTN, PRDX6, PROS1, PZP, RAP1B, S100A11, S100A6, SAA4, SERPINA1, SERPINA10, SERPINA3, SERPINE2, SERPINF2, SERPING1, SPP2, TALDO1, TKT, TPM1, TTR, TUBA1B, TUBB, UBA52, VCAN, VCP, VTN

**Male-only Biomarker Candidates**

ACTB, ACTG1, ALDH2, ANPEP, ANXA7, ARPC3, ASAH1, C1R, C4BPB, C8G, CALR, CANX, CAPG, CAPZB, CAST, CCT5, CNN3, COL14A1, COL1A1, COL5A2, CSPG2, DPYSL2, DSTN, EEF1A1, EEF1D, EHD2, EMILIN1, FBLN2, GAA, GANAB, GNB1, GPI, H3C15, HADHA, HAPLN1, HBB, HDGF, HNRNPA2B1, HNRNPD, HNRNPK, HSPA9, IGH@, IGK, ILK, ITGA2B, ITIH3, KLKB1, KRT9, LDHB, LTF, MAP4, MARCKS, MMP9, MPO, ORM2, PALLD, PCOLCE2, PCYOX1, PDCD6IP, PDIA4, PDIA6, PDLIM3, PDLIM7, PGAM1, PGLYRP2, PLG, PPIB, PRDX2, PRKCSH, PTGIS, QSOX1, RPLP2, S100A9, SAMHD1, SEPTIN7, SERBP1, SERBP1, SERPINH1, SH3BGRL, SH3BGRL3, SH3BGRL3, SLC2A1, SLC4A1, SOD1, SPTAN1, SSB, STAB1, TGFB1I1, THBS2, TPI1, TPM2, TPM3, TPP1, TPT1, WDR1, YWHAE, YWHAQ

### Shared and unique biomarker candidates between ageing IVD and age-dependent atherosclerosis in artery

**Shared Biomarker Candidates between Ageing (IVD) and Age-dependent Atherosclerosis (Artery)**

A2M, ACTB, ACTN4, AHNAK, AMBP, ANXA7, APOA1, APOB, APOE, ARHGDIB, C2, C3, C6, C9, CALU, CAST, CDH13, CLTC, COL14A1, COL1A1, COL4A2, COL5A2, COL6A1, COL6A2, COL6A3, DPYSL2, EEF1A1, EMILIN1, F13A1, F2, FBLN1, FLNA, FN1, GAA, GANAB, GAPDH, H4C1, HNRNPK, HSP90AB1, HSPA1A, HSPA1B, HSPG2, HTRA1, ITGB1, ITIH4, KNG1, KRT9, LAMC1, LRP1, LTBP2, MYH10, MYH11, NID1, ORM2, PCOLCE2, PDCD6IP, PDIA6, PFN1, PGD, PLEC, PLG, PLS3, PLTP, PRKCSH, RAP1B, S100A9, SERPINA1, SERPINA3, SERPINA4, SERPIND1, SERPING1, SLC4A1, SOD3, SPTA1, TF, THBS1, THBS2, TIMP1, TKT, TNC, TNXB, TTR, TUBA1B, VCAN, VTN

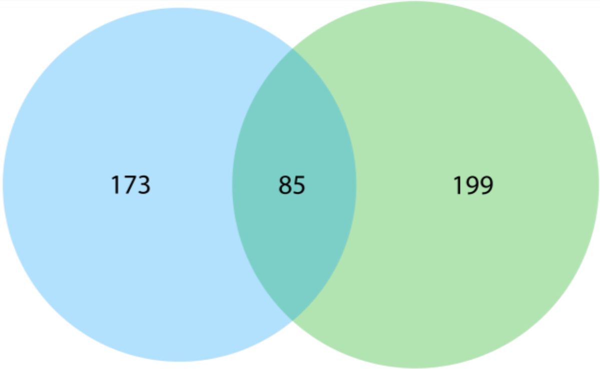


**Atherosclerotic Artery-only Biomarker Candidates**

ACTC1, ANK1, ANXA1, AOC3, APOL1, APOM, AQP1, C4B, C7, C8B, CAVIN1, CFI, COL12A1, COL18A1, CSRP1, CTSB, DPYSL3, EEF2, FCN3, FERMT3, FMOD, FTH1, GM10881, GPNMB, GSTO1, IQGAP1, LAMP1, LASP1, LBP, MMP12, MYH9, OGN, RBP4, RTN4, SPTB, TLN1, YWHAG, ACTG1, ALDH2, ANPEP, ARPC3, ASAH1, C1R, C4BPB, C8G, CALR, CANX, CAPG, CAPZB, CCT5, CNN3, CSPG2, DSTN, EEF1D, EHD2, FBLN2, GNB1, GPI, H3C15, HADHA, HAPLN1, HBB, HDGF, HNRNPA2B1, HNRNPD, HSPA9, IGH@, IGK, ILK, ITGA2B, ITIH3, KLKB1, LDHB, LTF, MAP4, MARCKS, MMP9, MPO, PALLD, PCYOX1, PDIA4, PDLIM3, PDLIM7, PGAM1, PGLYRP2, PPIB, PRDX2, PTGIS, QSOX1, RPLP2, SAMHD1, SEPTIN7, SERBP1, SERBP1, SERPINH1, SH3BGRL, SH3BGRL3, SH3BGRL3, SLC2A1, SOD1, SPTAN1, SSB, STAB1, TGFB1I1, TPI1, TPM2, TPM3, TPP1, TPT1, WDR1, YWHAE, YWHAQ, ACTR2, ACTR3, AEBP1, AFM, AGT, ARPC2, ASPN, C1QA, C1QB, CD14, CD36, CD44, CFHR1, CFL1, CLIC1, CNDP2, CNN1, COL1A2, COL4A1, CORO1A, CPN2, CRIP2, CYB5R3, DEFA1, DEFA3, EFEMP1, EIF4A1, EMILIN2, F10, F12, FBLN5, FGG, FLJ00385, GAS6, H3-3A, HPR, IGFALS, IGHG3, ITGB2, LTBP1, MYL12A, MYL9, PDIA3, PLD3, PON1, POSTN, PRDX6, PROS1, PZP, S100A11, S100A6, SAA4, SERPINA10, SERPINE2, SERPINF2, SPP2, TALDO1, TPM1, TUBB, UBA52, VCP

**Ageing IVD OAF-only Biomarker Candidates**

A1BG, ABI3BP, ACAN, ACLY, ACTN1, ACTN3, AKR1A1, ANGPTL2, ANGPTL5, ANXA2, ANXA5, APCS, APOA4, ARF1, ARF3, ATP1A1, BGN, C1S, C4A, CA2, CALD1, CALM1, CALM2, CALM3, CAPS, CAVIN3, CCDC80, CCN2, CD109, CDH1, CFH, CHAD, CHRDL2, CILP, CILP2, CKM, CLEC3A, CLEC3B, COL11A1, COL11A2, COL15A1, COL2A1, COL3A1, COL5A1, COL8A1, COL9A1, COL9A2, COMP, CPNE3, CPXM2, CRYAB, CSPG4, CTGF, CTSG, DF, DKK3, DPT, DYNC1H1, ECM1, ECM2, EEF1A1P5, EEF1G, EIF4A2, EMILIN3, ENO3, ENPP1, ENPP2, ERP44, EZR, FBN1, FBXO2, FGA, FGB, FHL1, FLNB, FLNC, FNDC1, FRZB, FSCN1, FTL, GBE1, GC, GCS1, GDI2, GLUD1, GSN, GSTP1, H1-0, H1-5, H2BC11, H2BC21, H2BC3, HAPLN3, HBD, HEXA, HHIPL2, HIST1H1E, HNRNPA3, HNRNPH1, HNRNPU, HP, HPX, HRG, HSPA5, HSPA8, HTRA3, IDH2, IGFBP7, IGHM, IGK@, IGL@, IGLC1, ISLR, ITIH1, ITIH2, ITIH6, KRT1, KRT10, KRT19, LAMA2, LAMA5, LAMB1, LAMB2, LCP1, LGALS3BP, LOX, LOXL3, LRG1, LTA4H, MAP1B, MATN2, MDH2, MELTF, MMP2, MOGS, MRC2, MSN, MXRA5, MYOM2, NACA, NCL, NID2, NNMT, NUCB1, NUCB2, OAF, OLFML1, ORM1, P4HA1, PAM, PCOLCE, PDGFC, PEBP1, PGK1, PLOD1, PPIC, PRDX3, PRG4, PRKAR1A, PRTN3, PSMA7, PTRF, PXDN, PXYLP1, PYGL, PYGM, RACK1, RBMX, RHOA, RPN1, RPN2, SCIN, SEMA3C, SEMA3E, SERPINB6, SERPINE1, SERPINF1, SET, SMOC1, SMOC2, SNC73, SPARC, SPTBN1, SRI, SRPX2, TGM2, THBS3, THBS4, TIMP3, TINAGL1, TUBA1A, TUBB4B, TXNDC5, UGDH, UGP2, VASN, VIM, VWA1, XYLT1
